# Microglial NF-κB drives tau spreading and toxicity in a mouse model of tauopathy

**DOI:** 10.1101/2021.02.22.432272

**Authors:** Chao Wang, Li Fan, Lihong Zhan, Lay Kodama, Bangyan Liu, Marcus Chin, Yaqiao Li, David Le, Yungui Zhou, Carlo Condello, Lea Grinberg, William W. Seeley, Bruce Miller, Sue-Ann Mok, Jason Gestwicki, Li Gan

**Affiliations:** Gladstone Institute of Neurological Disease, Department of Neurology, University of California, San Francisco, San Francisco, CA, USA; Helen and Robert Appel Alzheimer’s Disease Institute, Brain and Mind Research Institute, Weill Cornell Medicine, New York, NY, USA; Institute for Neurodegenerative Disease, Department of Pharmaceutical Chemistry, University of California, San Francisco, San Francisco, CA, USA; Institute for Neurodegenerative Disease, Department of Neurology, University of California, San Francisco, San Francisco, CA, USA; Memory and Aging Center, Department of Neurology, University of California, San Francisco, San Francisco, CA, USA

## Abstract

Activation of microglia, the brain’s innate immune cells, is a prominent pathological feature in tauopathies, including Alzheimer’s disease. How microglia activation contributes to tau toxicity remains largely unknown. Here we show that nuclear factor kappa-light-chain-enhancer of activated B cells (NF-κB) signaling, activated by tau, drives microglial-mediated tau propagation and toxicity. Constitutive activation of microglial NF-κB exacerbated, while inactivation diminished, tau seeding and spreading in PS19 mice, consistent with the observation that NF-κB activation accelerates processing of internalized tau fibrils in primary microglia. Remarkably, inhibition of microglial NF-κB specifically also rescued tau-mediated learning and memory deficits, and restored overall transcriptomic changes while increasing tau inclusions. On a single cell level, we discovered that tau-associated disease states in microglia were diminished by NF-κB inactivation and further transformed by constitutive NF-κB activation. Our study establishes a central role for microglial NF-κB signaling in mediating tau toxicity in tauopathy.

## Introduction

Abnormal aggregation and spreading of the microtubule-associated protein tau is the key defining feature of a group of heterogeneous neurodegenerative diseases known as tauopathies^1^. Alzheimer’s disease (AD) is the most common tauopathy, with a hallmark of neurofibrillary tangles (NFTs) composed of insoluble tau fibrils. Tau pathology correlates more closely with synaptic loss, neurodegeneration and cognitive decline than does amyloid pathology^2–4^. Understanding the pathogenic mechanism induced by pathological tau is critical for developing efficient therapeutics for AD and other tauopathies.

Neuroinflammation, the immune response in the central nervous system (CNS) characterized by reactive gliosis and increased inflammatory molecules, is one of the early and sustained pathological features of tauopathies^5^. Microglia, the resident innate immune cells in the CNS, are key players of neuroinflammation. Recent genetic and gene network analysis of late onsite AD (LOAD) identified several risk variants predominantly expressed in microglia, implicating a pivotal role of microglia in AD pathogenesis^6^. Reactive microglia have been observed to associate with NFTs in AD^7^, primary tauopathies^8, 9^ and tau transgenic models^10, 11^. *In vitro* studies confirm that tau directly activates microglia and triggers a pro-inflammatory profile^12^. Accumulated evidence also suggests that microglia are involved in tau-mediated pathobiology, including tau phagocytosis^13, 14^, post-translational modification and aggregation^15, 16^, spreading^17^ and tau-induced synaptic loss^18^. However, the molecular pathways underlying microglia-mediated tau toxicity remain poorly defined.

NF-κB is a transcription factor known to modulate many target genes that are associated with neuroinflammation, glial activation, oxidative stress, cell proliferation and apoptosis in the central nervous system. IκB kinase (IKK) activates NF-κB via phosphorylation and subsequent degradation of IκBα, an inhibitor of NF-κB ^19^. Genetic manipulations of IκB and IKK using the Cre-lox system enable conditional activation or inactivation of NF-κB and the dissection of functions of NF-κB in specific cell types^20^. Such studies have shown that neuronal NF-κB plays an essential role in synaptic plasticity and regulating learning and memory behaviors in basal conditions^21, 22^, whereas in pathological conditions, the anti-apoptotic role of neuronal NF-κB is associated with neuroprotective effects^23, 24^. Microglial NF-κB also regulates synaptic plasticity^25^.

Dysregulation of NF-κB has been implicated in AD pathogenesis^26, 27^. Indeed, in a metaanalysis, NF-κB signaling was found to be among the most perturbed pathways in LOAD brains ^28^. NF-κB is known to be activated by amyloid β (Aβ) and to contribute to Aβ production^29, 30^. We previously showed that specific inhibition of microglial NF-κB activation via deacetylation of p65 protected against Aβ toxicity in glial-neuron co-cultures^31^. However, very little is known about the role of microglial NF-κB activation in tauopathy.

Our current study establishes NF-κB signaling as a central transcription factor driving tau responses in *in vitro* and *in vivo* models of tauopathy. By genetically deleting or activating IKKβ kinase selectively in microglia, we investigated how microglial NF-κB signaling contributes to tau processing, seeding, and spreading, as well as tau toxicity using behavioral studies. Using bulk and single nuclei RNA-sequencing (RNA-seq), we dissected how microglial NF-κB activation and inactivation modify overall transcriptomic changes, tau-associated microglial states and underlying pathways in microglia. Our findings uncover novel mechanisms by which microglial NF-κB activation drives disease progression in tauopathy.

## Results

### Tau activates NF-κB pathway in microglia

To determine tau-induced transcriptome changes in microglia, we treated primary microglia with full-length wildtype (FL_WT) tau fibrils for 24 hours and analyzed them by RNA-seq. Synthetic recombinant tau monomers and fibrils contained negligible endotoxin (Supplementary table 1). Out of the 2975 differentially expressed genes (DEGs, False Discovery Rate (FDR)<0.05) (Fig. 1a and Supplementary table 2), Ingenuity Pathway Analysis (IPA) revealed that the top affected canonical pathways were associated with cellular immune responses, morphological changes, cell movement, survival and proliferation (Fig. 1b). NF-κB signaling, which is involved in the upstream or downstream regulation of many other canonical pathways, such as TNFR2, toll-like receptor (TLR), interferon and IL-12 signaling, was among the top altered cellular immune response pathways. Many NF-κB target genes were upregulated, including chemokines and receptors such as *Ccl5, Cxcl9,* complement component *C3*, proinflammatory cytokines *Il1b, Il12b,* and *Tnfa,* Fc fragment of IgG receptors involved in phagocytosis such as *Fcgr1,* and the NF-κB pathway component *Nfkbia* (Fig. 1a).

**Fig. 1.**
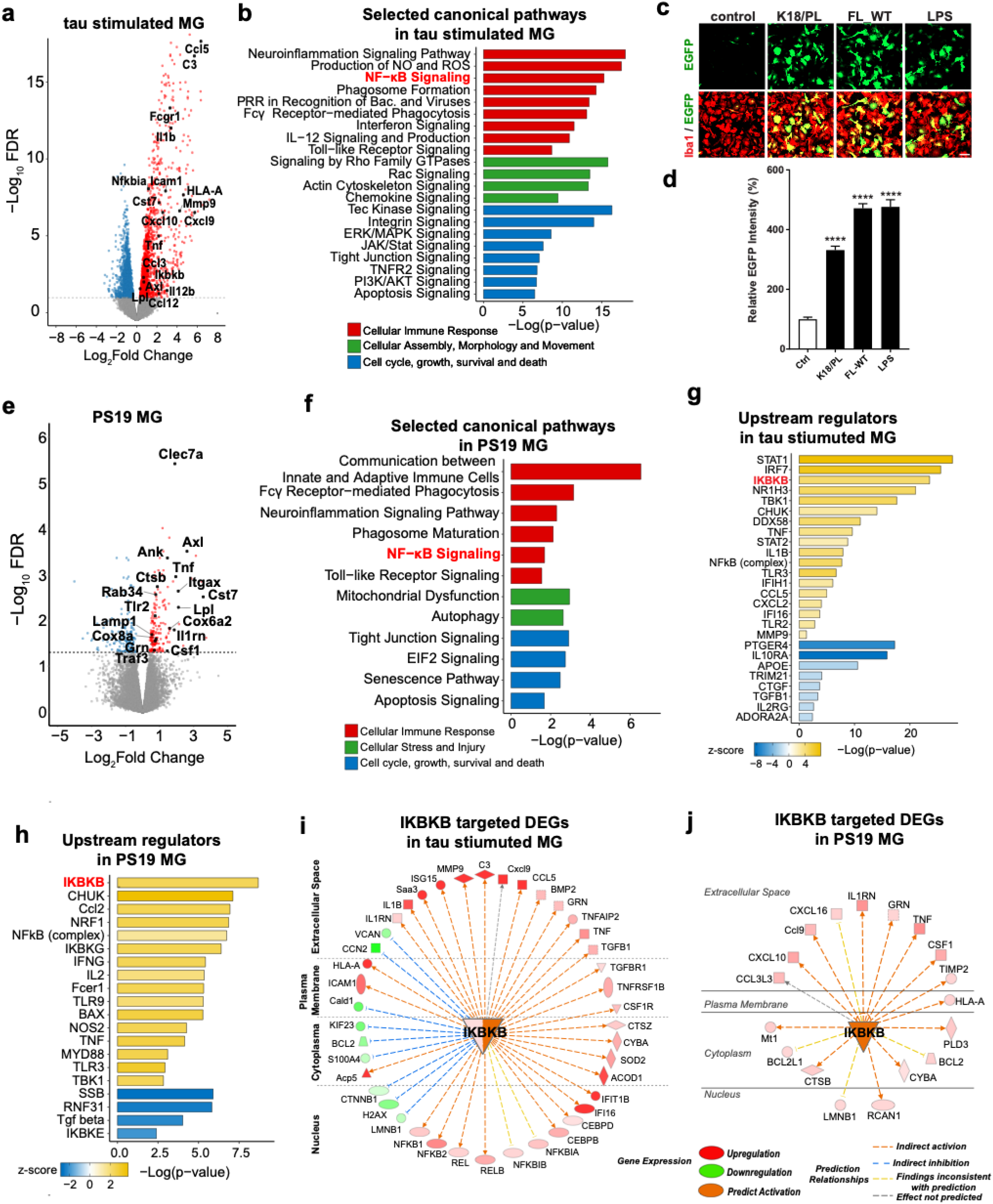
Tau activates NF-κB pathway in microglia. **(a)** Volcano plot of significant DEGs between full-length wildtype tau fibril-treated and vehicle-treated primary microglia. Colored points represent FDR<0.05, log_2_FC>0 upregulated genes (red) and log_2_FC<0 downregulated genes (blue). Selected NF-κB pathway associated genes and DAM signature genes are highlighted. FC, Fold change. **(b)** Selected IPA canonical pathways identified for significant DEGs of tau stimulated microglia. Canonical pathways are grouped by indicated categories. **(c,d)** Primary microglia infected with NF-κB reporter (Lenti-κB-dEGFP) virus were incubated with K18/PL fibrils (2.5ug/ml), full-length wildtype (FL-WT) tau fibrils (2ug/ml) and LPS (50ng/ml) for 24h. (**c**) Representative fluorescence high content images of EGFP (green) and microglial marker Iba1 (red); (**d**) Quantification of EGFP intensity. Scale bar, 50μm. Data are from three independent experiments, total N=21 wells. Values are mean ± SEM, relative to vehicle control, one-way ANOVA with Tukey’s multiple comparison post-test, ****p<0.0001 **(e)** Volcano plot of significant DEGs between isolated microglia from 11-month-old PS19 mice and non-transgenic controls. Selected NF-κB pathway associated genes and DAM signature genes are highlighted. **(f)** Selected IPA canonical pathways identified for significant DEGs in PS19 microglia. Canonical pathways are grouped by indicated categories. **(g,h)** Selected IPA predicted upstream regulators for DEGs of tau-stimulated microglia (**g**) and PS19 microglia (**h**). IKBKB (distinguished in red) is among the top upstream regulators. **(i,j)** The network of DEGs regulated by IKBKB in tau stimulated microglia (**i**) or PS19 microglia (**j**). Locations, gene expression levels and predicted relationships with IKBKB are illustrated as indicated labels. Shades of red and green represent log_2_FC of selected DEGs.

NF-κB transactivation was also measured with a reporter assay. Primary microglia were infected with lentivirus expressing EGFP under the control of the 5×κB enhancer element (Lenti-κB-dEGFP) ^31^. Both FL_WT and K18/PL tau fibrils, a truncated form of human tau fibrils containing only microtubule-binding domains with the P301L *MAPT* mutation^32^, significantly induced EGFP expression, indicating the activation of NF-κB promoter by tau fibrils (Fig. 1c, d). Wildtype or P301L mutant tau monomers also induced EGFP expression in microglia infected with Lenti-κB-dEGFP, confirming that both tau fibrils and monomers can activate microglial NF-κB pathway (Supplementary Fig. 1a,b). The extent of NF-κB activation induced by 2-2.5ug/ml tau fibrils was comparable to that of 50ng/ml lipopolysaccharide, which contains 250-2000 fold more endotoxin (Supplementary table 1).

We next profiled microglial transcriptional changes in a tauopathy mouse model of isolated microglia from 11-month-old PS19 mice. Disease-associated microglia (DAM) are identified as a subset of microglia associated with AD and other neurodegenerative diseases with a unique transcriptional signature^33, 34^. Some DAM signature genes, such as *Cst7, Axl, Lpl, Itgax, Clec7a, Cox6a2, Ank, Csf1,* and NF-κB target genes, such as *Tnfa, Il1rn, Tlr2,* and *Traf3,* were among the 187 upregulated DEGs (FDR<0.05, Supplementary Table 3) (Fig. 1e). NF-κB signaling was also identified as one of the top differentially regulated pathways in PS19 microglia (Fig. 1f). Other pathways and DEGs included cell growth and death, mitochondrial dysfunction and autophagy pathways, together with cytochrome-c oxidases, the terminal enzymes of the mitochondrial respiratory chain, such as *Cox6a2, Cox8a,* and endo-lysosome associated genes, such as *Ctsb, Ctss, Lamp1,Grn, Rab34,* all of which were upregulated in PS19 microglia (Fig 1e, f). Consistent with the canonical pathways associated with NF-κB activation, cell movement and migration, inflammatory responses and phagocytosis were the top activated biological functions in tau-stimulated microglia and PS19 microglia (Supplementary Fig. 1c, d). Specifically, IκB kinase complex subunit IKKβ *(Ikbkb)* was identified as one of the top upstream regulators responsible for tau-mediated transcriptomic changes in primary microglia (Fig. 1g) and PS19 microglia (Fig. 1h). *Ikbkb* gene itself was also upregulated in tau-stimulated microglia, and predicted to be activated in PS19 microglia (Fig. 1i, j). Indeed, a large repertoire of DEGs were predicted to be regulated by IKKβ, supporting IKKβ activation as a master regulator of tau-mediated microglial NF-κB activation.

### NF-κB transforms transcriptomes in cultured microglia

To directly investigate cell-autonomous effects of NF-κB in microglia, we activated IKKβ in microglia by crossing *Cx3cr1^CreERT2^* mice^35^ with R26-Stop^FL^ikk2ca mice, in which a constitutively active form of IKKβ was inserted into the Floxed-Rosa locus (hereafter referred to as *“IkbkbCA^F/F^*” mice)^36^. We treated primary microglia from *Cx3cr1^CreERT2^;IkbkbCA^F/F^* mice with 4-hydroxy tamoxifen^37^ to induce Cre expression (Fig. 2a). Elevation of *Ikbkb* was confirmed by RT-qPCR (Fig. 2b). We compared the transcriptomes induced by IKKβCA with those induced by tau stimulation, and observed 693 shared DEGs (556 upregulated and 137 downregulated) (Fig. 2c and Supplementary table 4), suggesting that tau-induced alterations may be partially mediated by NF-κB activation. IPA analyses of shared DEGs showed that in tau-stimulated microglia, enhanced cell proliferation, movement, phagocytosis, cytotoxicity and reduced cell death (Fig. 2d), as well as elevation of immune response pathways such as interferon, TLR signaling and inhibition of apoptosis signaling (Fig. 2e), may be mediated through NF-κB activation.

**Fig. 2.**
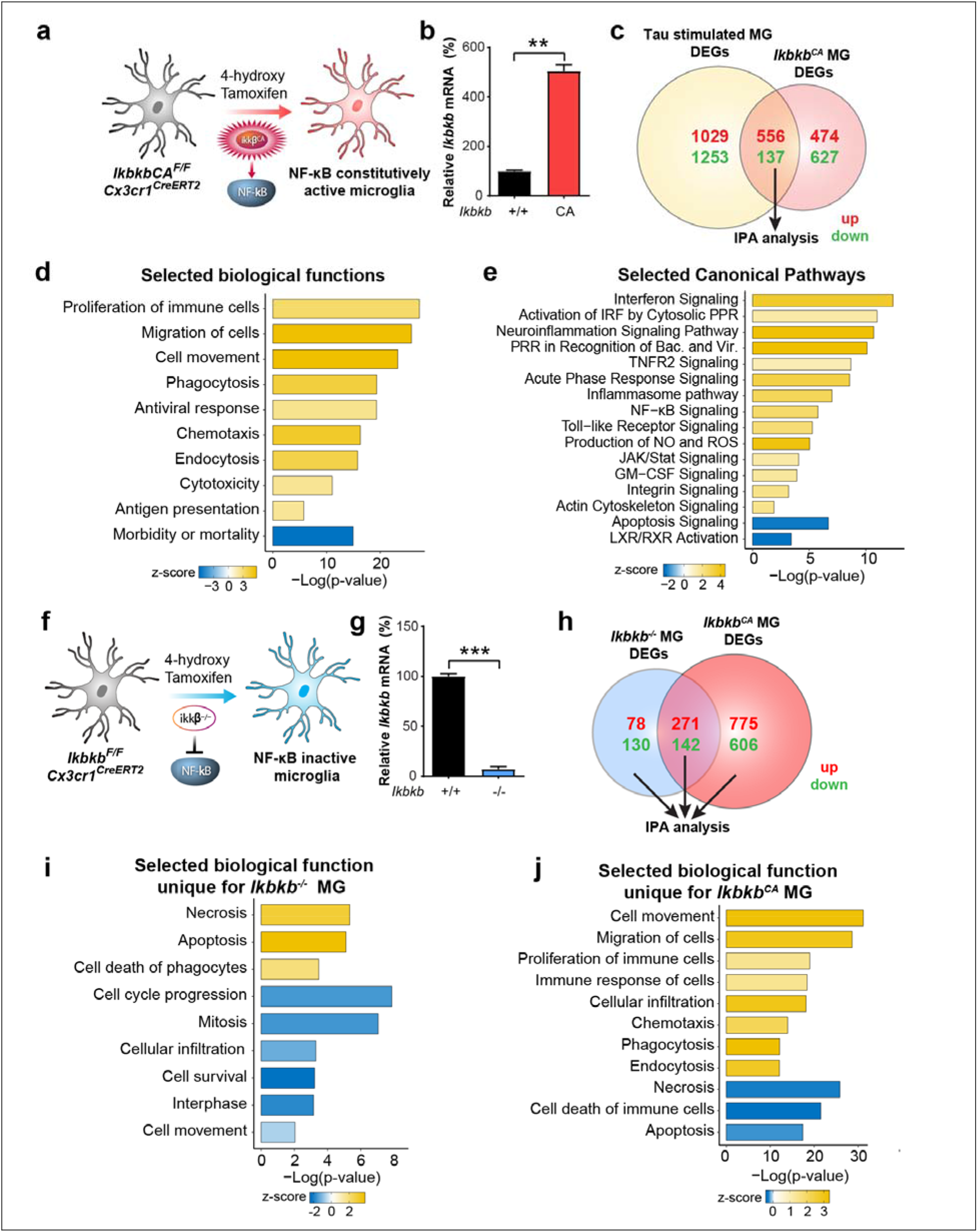
NF-κB dependent transcriptomic reprograming in primary microglia. **(a)** Experimental diagram illustrating generation of NF-κB constitutively active microglia. *IkbkbCA^F/F^ Cx3cr1^CreERT2^* primary microglia were incubated with 4-hydroxytamoxifen to induce IKKβCA expression. **(b)** Increased *Ikbkb* mRNA levels were confirmed by quantitative PCR analysis. N=3 per genotype. Values are mean ± SEM, relative to vehicle control. **P<0.01, unpaired student t-test **(c)** Venn diagram comparing DEGs between *Ikbkb^CA^* microglia and tau-stimulated microglia. DEGs: FDR < 0.05 in comparison to corresponding controls. Up-regulated gene numbers and *CA* down-regulated gene numbers are shown in red and green, respectively. DEGs shared by *Ikbkb^CA^* and tau-stimulated microglia are subjected to IPA analysis. **(d,e)** Selected IPA biological functions (**d**) and canonical pathways (**e**) identified for shared DEGs of *Ikbkb^CA^* microglia and tau-stimulated microglia identified in (**c**). **(f)** Experimental diagram illustrating generation of NF-κB inactivated microglia. *Ikkβ^F/F^ Cx3cr1^CreERT2^* primary microglia were incubated with 4-hydroxytamoxifen to delete IKK. **(g)** Decreased *Ikbkb* mRNA levels were confirmed by quantitative PCR analysis. N=3 per genotype. Values are mean ± SEM, relative to vehicle control. ***P<0.001, unpaired student t-test **(h)** Venn diagram comparing DEGs between *Ikbkb^-/-^* and *Ikbkb^CA^* microglia. DEGs: FDR < 0.05 in comparison to corresponding wildtype control, respectively. Unique and shared DEGs of *Ikbkb^-/-^* and *Ikbkb* microglia are subjected to IPA analysis. **(i, j)** Selected IPA biological functions identified for unique DEGs of *Ikbkb^-/-^* microglia (**i**) and *Ikbkb^CA^* microglia (**j**).

In complementary experiments, we selectively inactivated NF-κB in microglia by crossing *Cx3cr1^CreERT2^* mice with *Ikbkb^F/F^* mice ^38^, and treated primary microglia with 4-hydroxy tamoxifen^37^ to induce Cre expression (Fig. 2f). Deletion of *Ikbkb* in microglia was confirmed by RT-qPCR (Fig. 2g). To further examine NF-κB-dependent microglial transcriptomes, we next compared the DEGs between IKKβ null and IKKβCA microglia. Inactivation of NF-κB in microglia resulted in 208 unique DEGs (78 upregulated and 130 downregulated), while activation resulted in 1381 unique DEGs (775 upregulated and 606 downregulated) (Fig. 2h, Supplementary table 5). IPA analysis revealed that inactivation of microglial NF-κB led to pathways associated with elevated cell death, and decreased cell movement and proliferation. In direct contrast, activation of microglial NF-κB altered pathways associated with decreased cell death, but elevated cell movement, proliferation, and phagocytosis (Fig. 2i,j). Analyses of the top affected canonical pathways revealed that those unique for IKKβ null microglia were associated with cell cycle regulation, nucleotide biosynthesis, and DNA damage repair, whereas those unique for IKKβCA were enriched for integrin, TNFR2, and PI3K/Akt signaling (Supplementary Fig. 2a,b). Surprisingly, ~400 DEGs were shared by activating and inactivating microglial NF-κB (Fig. 2h and Supplementary table 6). Among the shared DEGs, a great fraction was involved in interferon signaling (Supplementary Fig. 2c). Specifically, interferon regulatory factor 3 & 7 *(Irf3, Irf7)*, interferon-α/β receptor *(Ifnar)* and interferon-γ *(Ifng)*, were the top upstream regulators predicted to be activated for these transcriptomic changes (Supplementary Fig. 2d). These results support the extensive cross-talk between interferons and the NF-κB pathway in microglia ^39^.

### NF-κB promotes tau processing in primary microglia

In cultured conditions, we found that Tau fibrils can be readily taken up by microglia, but not neurons, in a time-dependent manner, and once inside the cell they co-localize with the late endosome/lysosome labeled by Dextran^40^ (Supplementary Fig. 3a, 3b), where they can be proteolytically processed ^41, 42^. To determine the effects of NF-κB signaling on this process, we compared tau fibril accumulation in IKKβ null and IKKβCA microglia compared to their respective wildtype controls *(Ikbkb^+/+^* and *Ikbkb^WT^*). Inactivation of NF-κB enhanced, while activation diminished, the amount of tau fibrils remaining in microglia (Fig. 3a, b). To dissociate uptake from clearance, we performed a pulse-chase assay by preloading microglia with tau fibrils and assessing the time-dependent clearance in the next 24 hours (Fig. 3c). Compared with corresponding wildtype controls, inhibition of NF-κB slowed down, while activation accelerated, tau clearance in microglia (Fig. 3d-g). Thus, NF-κB activation in microglia reduces intracellular retention of tau by enhancing tau processing.

**Fig. 3.**
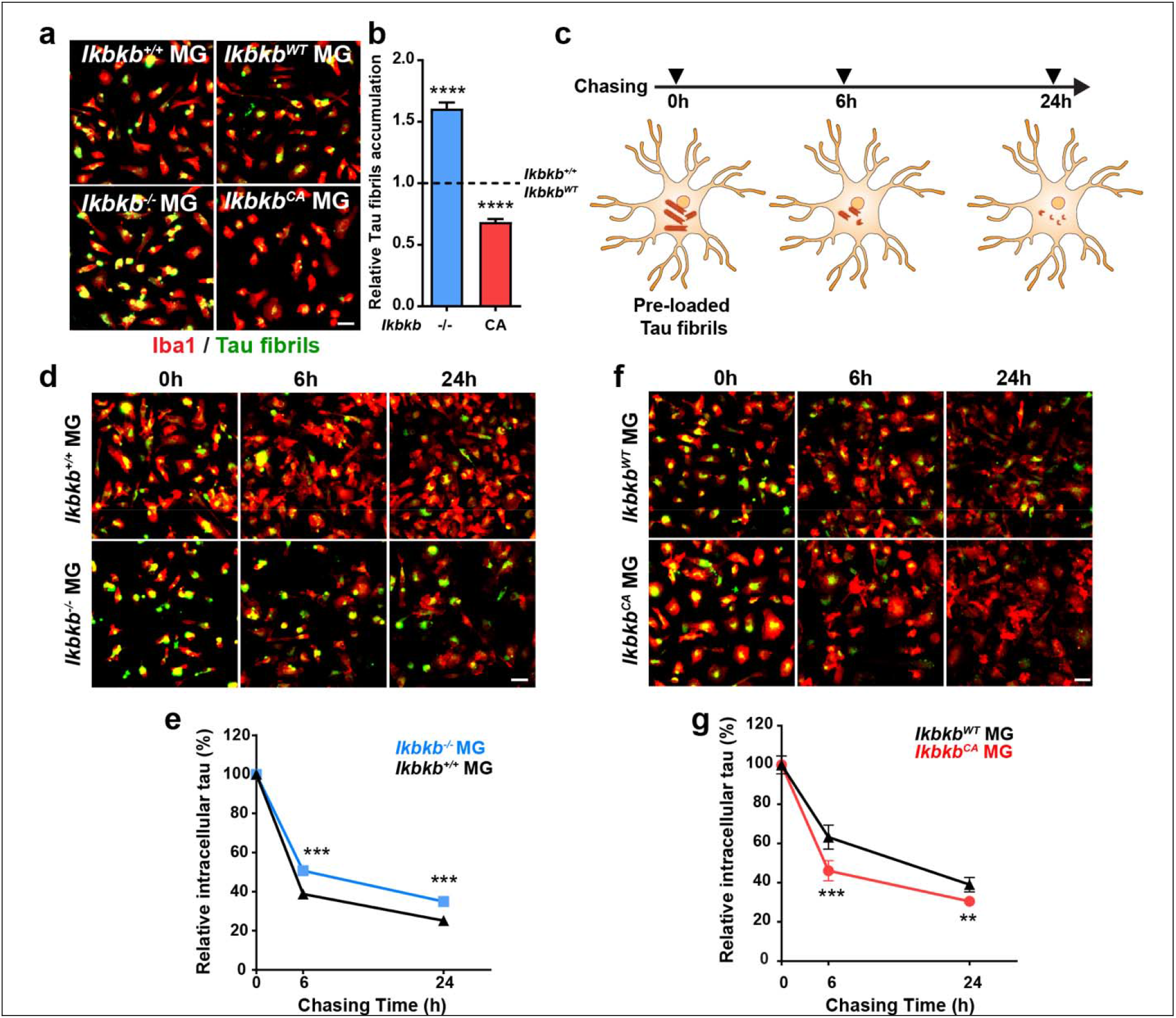
NF-κB promotes tau processing in primary microglia. **(a,b)** Representative high content fluorescence images **(a)** and quantification **(b)** of 24 hour fluorescent tau fibril accumulation in primary *Ikbkb^-/-^* and *Ikbkb^CA^* microglia compared with *Ikbkb^+/+^* and *Ikbkb^WT^* control, respectively. Scale bar, 25 μm. Data are from three independent experiments, total N=9 wells. values are mean ± SEM, relative to *Ikbkb^+/+^* and *Ikbkb^WT^* control, ****p<0.0001, one-way ANOVA with Tukey’s multiple comparison post-test. **(c)** Schematic diagram illustrating pulse chase assay designed for quantification of tau clearance in microglia. Primary microglia were pre-loaded with fluorescent tau fibrils for 24 hours, followed by chase for 6 hours and 24 hours. **(d-g)** Representative high content fluorescence images (**d,f**) and quantifications (**e,g**) of pulse chase assay in *Ikbkb^-/-^* microglia (**d,e**) and in *IkbkbCA* microglia (**f,g**). Scale bar, 25 μm. Data are from three independent experiments, total N=18 wells. values are mean ± SEM, relative to 0h pulse control, ***p<0.001, **p<0.01 STATA mixed-effects model.

### Microglial NF-κB activation promotes tau seeding and spreading in PS19 mice

Tau pathology spreads from entorhinal cortex to the hippocampal region in early stage of AD ^43^. Microglia were reported to contribute to this progress through secreting processed tau^17^. Seeding and spread of tau inclusions can be modeled by inoculating young PS19 mice with exogenous tau seeds ^32^ (Supplementary Fig. 4a). To investigate the role of microglia on tau seeding and spreading, we depleted microglia by feeding young PS19 mice a diet containing the colony stimulating factor 1 receptor inhibitor PLX5622 (PLX)^44^, followed by inoculating with either brain extract from progressive supranuclear palsy (PSP) patients (Supplementary Fig. 4b) or synthetic K18/PL tau fibrils (Fig. 4a) unilaterally into hippocampus. PSP-injected mice were continuously fed with PLX for three months to prevent microglial repopulation, which reduced the number of microglia over 80% (Supplementary Fig. 4c,d). Depletion of microglia significantly reduced the number of tau inclusions in AT8+ neurons in both contralateral and ipsilateral cortex, consistent with the notion that microglia promote tau seeding and spreading in mouse tauopathy models (Supplementary Fig. 4e,f). Similarly, microglial depletion significantly reduced the seeding and spreading of tau in PS19 mice inoculated with synthetic K18/PL tau fibrils, which induced tau spreading within 1 month (Fig. 4a-c). Importantly, no AT8+ or MC1 + neurons were detected in PS19 mice inoculated with PBS, or in non-transgenic control mice inoculated with tau seeds (Supplementary Fig. 4g).

**Fig. 4.**
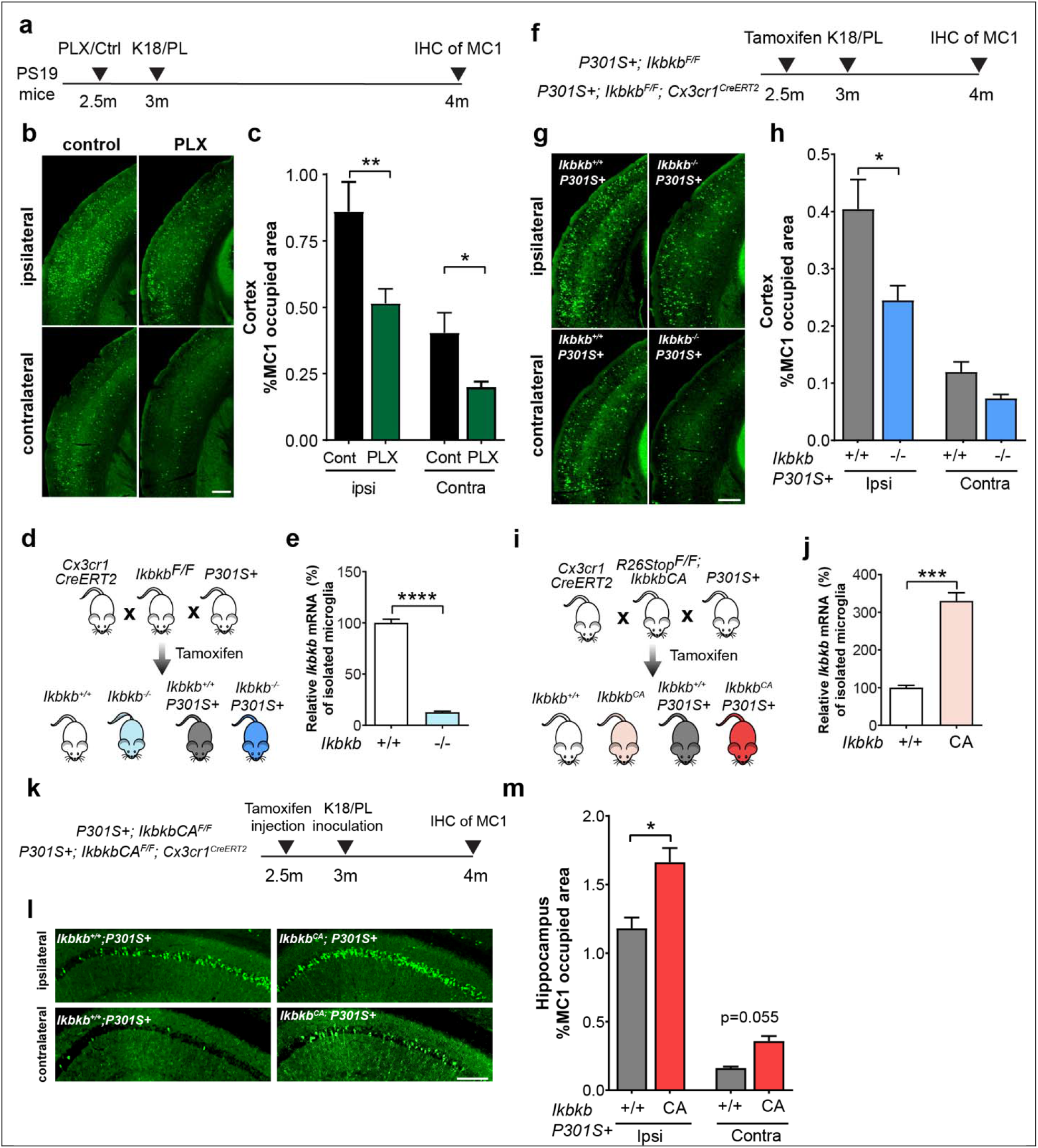
Microglial NF-κB activation promotes tau seeding and spread in PS19 mice. **(a)** Schematic diagram illustrating the experimental design and timeline of microglia depletion and K18/PL tau fibril inoculation in PS19 mice. Pathological tau seeding and spreading was determined by immunohistochemical staining of MC1. **(b,c)** Representative immunohistochemical staining of MC1 tau positive neurons in ipsilateral and contralateral cortex (**b**) and quantification of percentage of MC1 occupied area (**c**) in control diet feeding mice (n=6) and PLX diet feeding mice (n=8), 6 sections per mice. ** p<0.01, * p<0.05, STATA mixed model. Scale bar, 500μm **(d)** Breeding diagram illustrating the generation of PS19 mice with microglia-conditional deletion of IKKβ. **(e)** Quantitive PCR analysis shows a dramatic reduction of *Ikbkb* mRNA level in adult microglia isolated from *Ikbkb^-/-^* mice (n=5) compared with wildtype control *Ikbkb^+/+^* (n=3) mice. ****P<0.0001, unpaired student t-test. **(f)** Schematic diagram illustrating the experimental design and timeline of microglial IKKβ deletion and K18/PL tau fibrils\ inoculation in PS19 mice. **(g,h)** Representative immunohistochemical staining of MC1 tau positive neurons in ipsilateral and contralateral cortex (**g**) and quantification of the percentage of MC1 occupied area (**h**) in *Ikbkb^+/+^;P301S+* (n=10) and *Ikbkb^-/-^;P301S+* (n=15)mice. 5 sections per mouse. *P<0.05, STATA mixed-effects model. Scale bar, 500μm. **(i)** Breeding diagram illustrating the generation of PS19 mice with microglia expressing constitutively active form of IKKβ. **(j)** Quantitive PCR analysis shows a dramatic increase of *Ikbkb* mRNA level in adult microglia isolated from *Ikbkb^CA^* mice (n=4) compared with wildtype control *Ikbkb^+/+^* mice (n=3). ***P<0.001, unpaired student t-test. **(k)** Schematic diagram illustrating the experimental design and timeline of expressing microglial IKKβCA and K18/PL tau fibril inoculation in PS19 mice. **(l,m)** Representative immunohistochemical staining of MC1 tau positive neurons in ipsilateral and contralateral hippocampus CA1 region (**l**) and quantification of the percentage of MC1 occupied area (**m**) in *Ikbkb^+/+^;P301S+* (n=9) and *Ikbkb^CA^;P301S+* (n=14) mice. 8 sections per mouse. *P<0.05, STATA mixed-effects model. Scale bar, 200μm

Since microglial processing of tau is regulated by NF-κB, we reasoned that microglial NF-κB activity could also affect tau seeding and spreading *in vivo.* We selectively deleted *Ikbkb* in adult microglia of PS19 mice by crossing *Cx3cr1^CreERT2^* with *Ikbkb^F/F^* and PS19 mice. Tamoxifen injection diminished IKKβ expression in adult microglia from *Cx3cr1^CreERT2^; Ikbkb^F/F^* (referred to as *Ikbkb^-/-^)* mice, as confirmed with qRT-PCR analyses (Fig. 4d,e). To measure how tau seeding and spreading were affected, 3-month-old mice were inoculated with tau fibrils in one-side of hippocampus two weeks following tamoxifen injection (Fig. 4f). K18/PL fibrils were used to induce tau spreading since only 4 week post-inoculation time is needed for robust widespread tau propagation in the cortex. Inactivation of microglial NF-κB reduced the amount of MC1+ tau inclusions significantly at the ipsilateral side of the cortex (Fig 4g,h), similar to the effect of depleting microglia (Fig. 4c), suggesting a critical role for microglial NF-κB activation in tau spreading. In complementary experiments, tamoxifen injection enhanced IKKβ expression in adult microglia from *Cx3cr1^CreERT2^;IkbkbCA^F/F^* (referred to as *ikbkb^CA^*) mice (Fig. 4i,j). We reasoned activating microglial NF-κB could enhance tau propagation. To avoid ceiling effects, we reduced the amount of inoculated tau fibrils than that in microglial-depletion experiments. One-month after the inoculation, *ikbkb^CA^* mice exhibited elevated tau inclusions significantly at the ipsilateral side, with modest increase at the contralateral side of hippocampus (Fig. 4k-m). Our findings indicate that NF-κB activation is required to promote microglial-mediated tau seeding *in vivo.*

### Inactivation of microglial NF-κB partially restores microglia homeostasis and protects against spatial learning and memory deficits in PS19 mice

Microgliosis and amoeboid morphological changes are early and sustained phenomena in PS19 mice^10^. Inactivation of microglial NF-κB reduced microgliosis in the hippocampus of 8 to 9-month-old PS19 mice (Fig. 5a,b), and a similar trend was observed in the cortex (Fig. 5c,d). Imaris analysis of microglial morphology further revealed that inhibition of microglial NF-κB resulted in more ramified morphology with longer processes and more branches, partially reversing the amoeboid morphology induced by pathogenic tau (Fig. 5e–g).

**Fig. 5.**
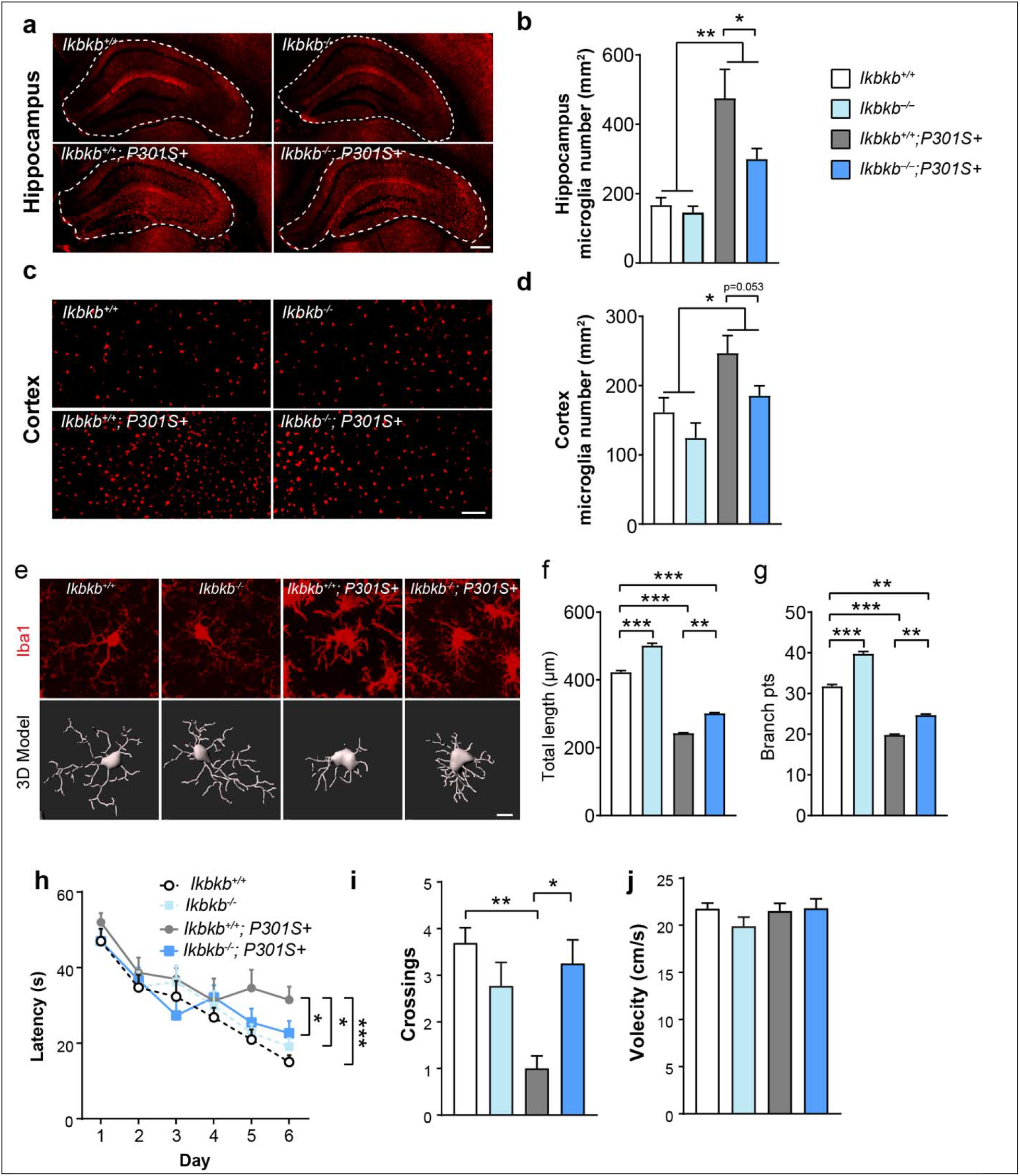
Inactivation of microglial NF-kB partially rescues microgliosis, morphological alterations and protects against spatial learning and memory deficits in PS19 mice. **(a-d)** Representative immunohistochemical staining **(a,c)** and quantification **(b,d)** of Iba1+ microglia in the hippocampus (**a,b**) and cortex (**c,d**) of *Ikbkb^+/+^* (n=3), *Ikbkb^-/-^*(n=3), *Ikbkb^+/+^;P301S+* (n=9), *and Ikbkb^-/-^;P301S+* (n=12) mice. Scale bar, hippocampus 250μm, cortex 100 μm. **P < 0.01, *P < 0.05, Two-way ANOVA with Sidak’s multiple comparisons post-test **(e)** Representative immunohistochemical confocal images showing morphological features of microglia from the hippocampus of *Ikbkb^+/+^*, *Ikbkb^-/-^, Ikbkb^+/+^;P301S+, and Ikbkb^-/-^;P301S+* mice and the corresponding 3D reconstructions using Imaris. Scale bar, 10μm **(f,g)** Quantification of the total length of microglial processes (**f**) and the number of microglia process branch points (**g**). N=4 mice per genotype (4 sections per mouse) were imaged and >250 Iba1+ microglia from each mouse were analyzed. **P < 0.01, ***P < 0.001, STATA mixed-effects model. **(h-j)** Morris Water Maze test was performed using *Ikbkb^+/+^* (n=13), *Ikbkb^-/-^*(n=13), *Ikbkb^+/+^;P301S+* (n=8), *and Ikbkb^-/-^;P301S+* (n=12) mice.(**h**) Escape latency was plotted against the training days; **(i)** Times of crossing the platform location in the 72-hour probe trial; **(j)** swimming speed ***p<0.001, ** p<0.01,*p<0.05 Two-way ANOVA with Tukey’s multiple comparisons post-test.

PS19 mice exhibit spatial learning and memory deficits starting at 7–8 months of age^45, 46^. To examine the functional outcome of inhibition of microglial NF-κB activity in PS19 mice, we tested 8 to 9-month-old *Ikbkb^+/+^, Ikbkb^-/-^, Ikbkb^+/+^;P301S+ and Ikbkb^-/-^;P301S+* mice in the Morris water maze (MWM) test, a hippocampus-dependent assay that evaluates spatial learning and memory deficits. Inhibition of microglial NF-κB alone did not significantly impact spatial learning and memory, measured by the learning curve and number of times to cross platform location in a 72-hour probe trial (Fig. 5h,i). Strikingly, inactivating microglial NF-κB in PS19 mice significantly improved the learning ability (Fig. 5h) and restored the spatial memory in the probe trial (Fig. 5i), without affecting swimming speed (Fig. 5j). Thus, hyperactive microglial NF-κB plays a critical role in altering microglial homeostasis and driving cognitive deficits in PS19 mice.

### Inactivation of microglial NF-κB rescues tau-mediated transcriptomic changes while increasing tau inclusions

We next examined if the protective effects of microglial NF-κB inactivation are mediated by reducing tau inclusions. Surprisingly, instead of reducing neuronal tau inclusions, inactivation of microglial NF-κB markedly increase the number of intraneuronal tau inclusions in both hippocampus and cortex of 9–10 month old PS19 mice (Fig. 6a-c). This indicates that neuronal tau inclusions might not be toxic per se if maladaptive microglial responses are blocked by inactivation of NF-κB.

**Fig. 6.**
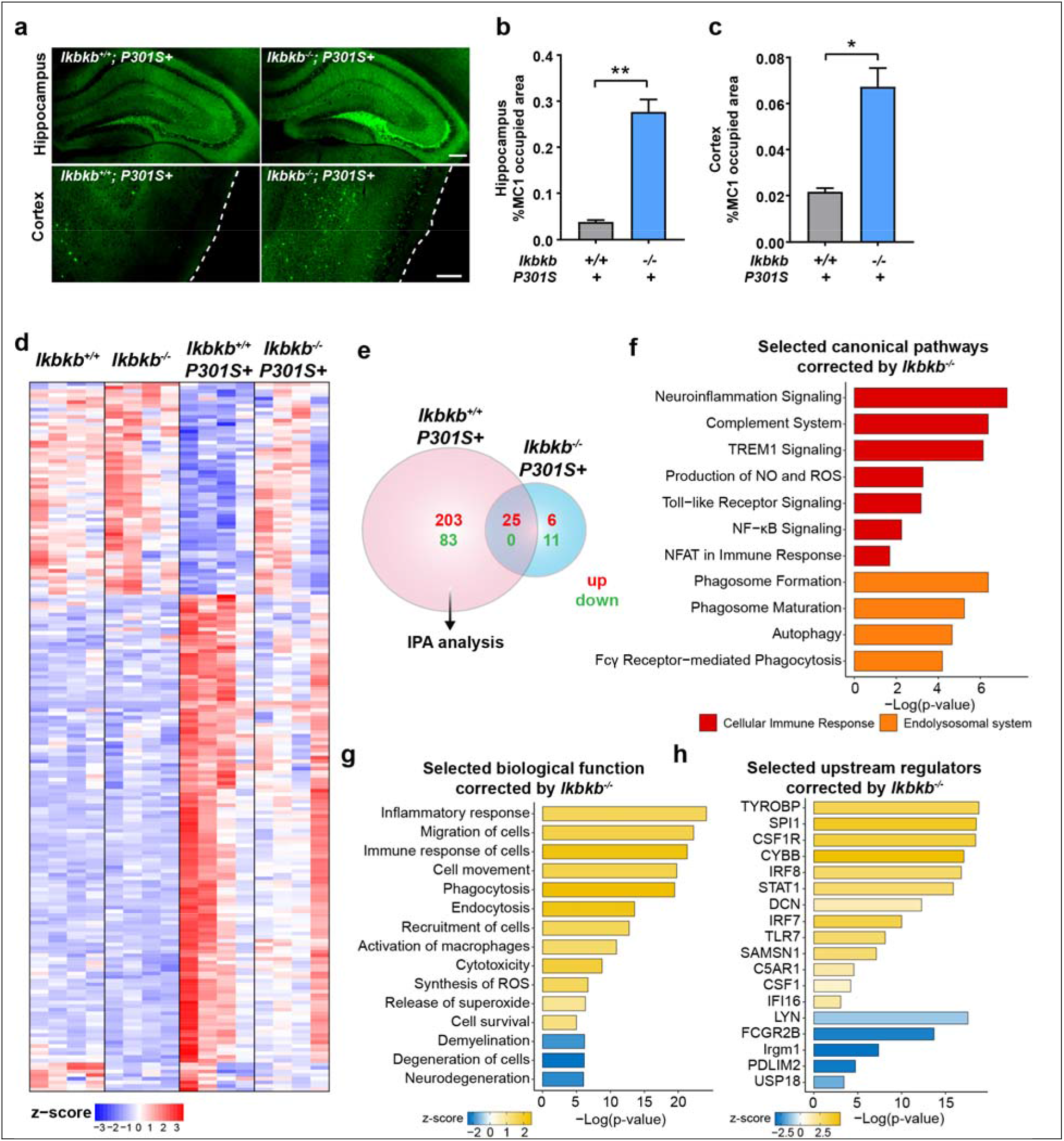
Inactivation of microglial NF-kB rescued tau-mediated transcriptomic changes while increasing tau inclusions. **(a)** Representative immunohistochemical staining of MC1 tau in the hippocampus and cortex of *Ikbkb^+/+^;P301S+* mice and *Ikbkb^-/-^;P301S+* mice. Scale bar, 250μm **(b,c)** Quantification of the percentage of MC1occupied area in the hippocampus (**b**) and cortex (**c**) of *Ikbkb^+/+^;P301S+,* n=9 mice; *Ikbkb^-/-^;P301S+,* n=12 mice; 7-9 sections per mouse; *P < 0.05, ***P < 0.001, STATA mixed-effects model. **(d-h)** Cortical tissues from *Ikbkb^+/+^, Ikbkb^-/-^, Ikbkb^+/+^;P301S+ and Ikbkb^-/-^;P301S+* mice (n=4 mice/genotype) were used for bulk RNA-seq analysis. DEGs (FDR<0.05) of each genotype were generated in comparison to *Ikbkb^+/+^* mice. **(d)** Heat map representing significant DEGs (FDR<0.05) across all genotypes. Heatmap shows the value of log_2_PFKM of each gene, which was normalized and z-scored (color-coded). Shades of red represent up-regulation, and shades of blue represent down-regulation. DEGs and biological replicate samples were hierarchically clustered. **(e)** Venn diagram comparing the number and overlap of DEGs in *Ikbkb^+/+^;P301S+* and *Ikbkb^-/-^;P301S+* mice. 286 DEGs corrected by IKKβ deletion are subjected to IPA analysis. **(f-h)** Selected IPA canonical pathways (**f**), biological functions (**g**), and top predicted upstream regulators (**h**) identified for 286 DEGs corrected by microglial IKKβ deletion in PS19 mice that are shown in (**e**).

To dissect the mechanisms underlying the protective effects of inhibiting microglial NF-κB activity, we performed bulk RNA-seq of cortical tissues from 8 to 9-month-old *Ikbkb^+/+^, Ikbkb^-/-^, Ikbkb^+/+^;P301S+* and *Ikbkb^-/-^;P301S+* mice. Remarkably, inactivation of microglial NF-κB resulted in a reversal of more than 90% of DEGs (286 out of 311 genes) in PS19 mice (Fig. 6d, e and Supplementary table 7), despite elevating the tau inclusions. IPA analysis of these 286 genes revealed that the majority of reversed canonical pathways were related to inflammatory responses and the endo-lysosome system, and included neuroinflammation signaling, complement system, ROS generation, TLR signaling, and phagocytosis (Fig. 6f). Tau-mediated biological function alterations including inflammatory response, cell movement, phagocytosis, superoxide production, and demyelination were also reversed (Fig. 6g). The top upstream regulators of the pathways and functions reversed by NF-κB inactivation include those regulating immune cell survival and activation (e.g. *Tryobp, Spi1, Csf1r, Csf1)*, superoxide generation (e.g. *Cybb*), and the interferon pathway (e.g. *Irf7, Irf8, Ifi16, Stat1*), suggesting these pathways are likely to be associated with the toxic effects of microglia in PS19 mice (Fig. 6h).

### NF-κB is required for tau-associated microglial states in PS19 mice

Microglia exhibit disease-associate states in the presence of pathology^34^, including tau^47^, which have been characterized by single cell RNA-seq. To characterize the effects of NF-κB on microglial transcriptomes, we performed single nuclei RNA-seq (snRNA-seq) using cortical tissues from *Ikbkb^+/+^;P301S+, Ikbkb^-/-^;P301S+* and *Ikbkb^CA^;P301S+* mice. Non-transgenic *Ikbkb^+/+^* mice were used as non-disease controls. Following an established snRNA-seq protocol^48^, we sequenced 114,118 nuclei from all four genotypes. After removal of potential multiplets using DoubletFinder^49^ and filtering for low-quality nuclei determined by thresholding gene counts, UMI counts and percentage mitochondrial genes per nuclei (Supplementary Fig. 5a-e), we used 103,681 nuclei for downstream analysis. Using reference gene markers for annotations (Supplementary Fig. 5g), we identified major cell types of the brain which were similarly represented within each group and individual mouse (Fig. 7a, b, Supplementary Fig. 5f).

**Fig. 7.**
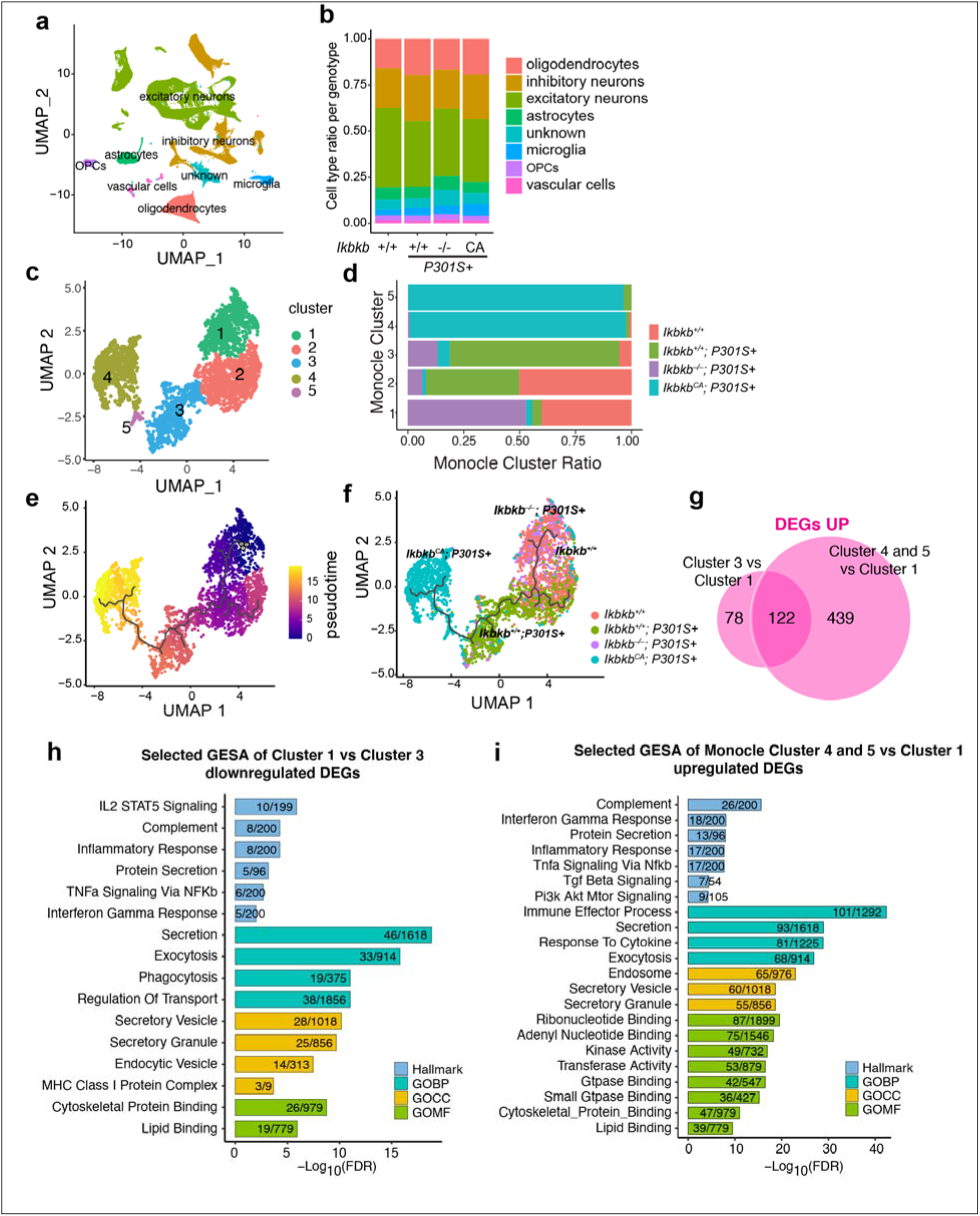
NF-κB is required to induce tau-associated microglia states in PS19 mice. Single nuclei RNA-seq were performed using cortical tissues from *Ikbkb^+/+^* (n=4), *Ikbkb^+/+^;P301S+*(n=3), *Ikbkb^-/-^;P301S+* (n=2) and *Ikbkb^CA^;P301S+* (n=2) mice. **(a)** UMAP plots of all single nuclei and their annotated cell types. OPCs: Oligodendrocyte progenitor cells. **(b)** Proportion of cell types for each genotype. **(c)** UMAP plot depicting different microglial cell sub-clusters. Each cell was color coded based on their cluster affiliation. **(d)** Proportion of cells in each microglial sub-cluster for each genotype. **(e)** Pseudotime-time trajectory demonstrating the shift of microglial state. **(f)** Pseudotime-time trajectory of microglia labeled with genotypes. The trajectory illustrates the shift of microglial transcriptome from *Ikbkb^-/-^;P301S+* to *Ikbkb^CA^;P301S+* mice. **(g)** Venn diagram comparing the number and overlap of downregulated DEGs in Cluster 1 vs Cluster 3, and Cluster 4 and 5 vs Cluster 1. (**h,i)** Selected GESA hallmark and Gene Ontology pathways identified for downregulated microglia DEGs in Cluster 1 vs Cluster 3 (**h**) and upregulated microglia DEGs in Cluster 4 and 5 vs Cluster 1 (**i**). GOBP: GO biological processes; GOCC: GO cellular compartment; GOMF: GO molecular function.

We then examined the trajectory of the subclusters of a total 4,498 microglia to investigate how NF-κB affects microglial states in P301S mice using Monocle^50^. The microglial population from the 4 genotypes exhibited a clearly-defined five subclusters (Fig. 7c). Analyses of the distribution of the five subclusters by genotypes revealed that microglia from *Ikbkb^+/+^* brains were distributed among clusters 1 and 2, while the vast majority of cluster 3 came from *Ikbkb^+/+^;P301S,* likely representing tau-induced microglial states (Fig. 7d). Most microglial from clusters 4 and 5 came from *Ikbkb^CA^; P301S*, suggesting that constitutive NF-κB activation further transforms microglial states (Fig. 7d). In direct contrast, microglia from *Ikbkb^-/-^;P301S* were mostly found in Cluster 1, along with wildtype *Ikbkb^+/+^* microglia, demonstrating that activation of NF-kB is required for tau-induced transcriptomal changes (Fig. 7d). We next examined microglial trajectory with pseudotime from 0 to 15 (Fig, 7e). Microglia from non-transgenic control (*Ikbkb^+/+^*) mice were enriched at the starting point of the trajectory (pink, Fig, 7e, f). Microglia from PS19 mice with wildtype IKKβ *(Ikbkb^+/+^;P301S+*) exhibit trajectory away from those of non-transgenic control (green, Fig. 7f), while those lacking IKKβ *(Ikbkb^-/-^;P301S+)* partially overlaps with that of *Ikbkb^+/+^* control, illustrating their failure to advance in the direction of that of PS19 microglia (purple, Fig. 7e). The other end of the trajectory (yellow, Fig. 7e) was populated with microglia from *Ikbkb^CA^;P301S+* mice, extending beyond the tau-associated disease states (cyan, Fig. 7f). Thus, tau-associated disease states in microglia requires NF-κB activation.

We further dissected the pathways modified by microglial NF-κB in PS19 mice. We identified 961 DEGs in *Ikbkb^-/-^;P301S+* microglia and 691 DEGs in *Ikbkb^CA^;P301S+* microglia (vs. *Ikbkb^+/+^; P301S+)* (FDR<0.05, |log2FC|>0.1, Supplementary Fig. 6a, b, Supplementary table 8). We found that genes induced by tau positively correlated with DAM genes (Supplementary Fig. 6c), while those induced by *Ikbkb^CA^* exhibited no correlation, indicating distinct microglial states (Supplementary Fig. 6d). Indeed, the DEGs upregulated in tau-induced cluster 3 vs. cluster 1 *(Ikbkb^-/-^;P301S+* and *Ikbkb^+/+^*) partially overlap with those upregulated by constitutive NF-κB activation, with 439 DEGs uniquely upregulated in *Ikbkb^CA^;P301S+* microglia (vs. *Ikbkb^+/+^; P301S+)* (FDR<0.05, |log2FC|>0.1, Fig. 7g, Supplementary table 9). Given the protective effects of NF-κB inactivation against tau-mediated toxicity, we were particularly interested in the 200 DEGs downregulated by NF-κβ inactivation by comparing gene expression in cluster 1 vs. cluster 3 (Fig. 7h), which could underlie its protective effects against functional deficits and tau seeding/spread. Investigation of the overlapping pathways downregulated by NF-κ? inactivation using gene set enrichment analysis (GSEA) identified complement, the IL2/STAT5, and lipid binding pathways (Fig. 7h, Supplementary Fig. 6a). Consistent with our finding that microglial NF-κB promotes tau processing, seeding and spread, both proteolysis and exocytosis functions of microglia were downregulated in *Ikbkb^-/-^;P301S+* microglia and upregulated in *Ikbkb^CA^;P301S+* microglia by comparing gene expression in clusters 4 and 5 vs. cluster 1 (Fig. 7i). These pathways include secretory granule complex, vesicular/protein transport and exocytosis, trafficking (Fig. 7h,i,). The reprogramming of microglial disease states by NF-κB inactivation provides molecular underpinnings for microglial-mediated tau seeding and tau-mediated cognitive deficits.

## Discussion

Here, we show that microglial NF-κB activation is required for microglial-mediated tau spreading and tau-mediated spatial learning and memory deficits in tauopathy mice. NF-κB signaling was among the top altered cellular immune response pathways in response to tau in microglia isolated from PS19 tauopathy mice. By genetically activating or inactivating microglial NF-κB, we found that NF-κB accelerates microglial processing of tau in cultured microglia, and promotes tau seeding and spreading *in vivo.* Moreover, inactivation of NF-κB in tauopathy mice partially restored microglial homeostasis, reversed tau-mediated transcriptomic changes, and rescued spatial learning and memory deficits. By identifying tau-associated microglial states that were diminished by NF-κB inactivation, our snRNA-seq analyses further reveal potential molecular mechanisms underlying microglial-mediated tau seeding/spreading and toxicity.

In both tau-stimulated primary microglia and isolated microglia from aged PS19 mice, we found that NF-κB pathway is among the top upstream regulators, consistent with a previous study of microglia from rTg4510 mice ^51^. In contrast, NF-κB target genes were not enriched in a microglial transcriptome study from APPswe/PS1dE9 mice, an Aβ driven AD model^52^, suggesting distinct disease-associated transcriptional programs induced by tau vs. Aβ. Microglial NF-κB could be activated by soluble tau monomers, in agreement with a previous study showing that microglial NF-κB is activated as early as 2 months of age in the rTg4510 model ^51^. We observed that some of the pathways and functions induced by tau overlapped with those induced by NF-κB activation, including proliferation and cell movement/migration, and that these could be reversed by inactivating microglial NF-κB *in vitro* and in PS19 mice. Moreover, inhibition of microglial NF-κB activity in PS19 mice reduced microgliosis, and resulted in longer and more branchy processes, partially reverting cells to a more homeostatic microglial state. Our snRNAseq analyses further revealed that NF-κB is required for tau-associated microglial states in PS19 mice. NF-κB inactivation diminished, while constitutive NF-κB activation extended beyond, the tau-associated disease states.

Emerging evidence supports the hypothesis that microglia participate in tau seeding and spreading^14, 17^. NLRP3–ASC inflammasome activation was found to exacerbate exogenously seeded tau pathology as well as non-exogenously seeded intraneuronal tau aggregates, at least partially through modulating tau phosphorylation^53, 54^. Our study shows that in PS19 mice, exogenous tau inoculation-induced tau seeding was accelerated by microglial NF-κB activation, but diminished by NF-κB inactivation. Indeed, inactivation of microglial NF-κB signaling alone induced a similar reduction in seeding as depleting microglia altogether, highlighting the central role of NF-κB signaling in microglia-mediated tau seeding. NF-κB activation could promote microglia to secrete more seeding-competent tau, and thus accelerate the spread of tau pathology. Consistent with this notion, we showed that microglial NF-κB activation accelerated the processing of tau, resulting in reduced intracellular tau retention in cultured microglia. Moreover, our snRNA-seq analysis of microglia in PS19 mice revealed that protein degradation and secretion are diminished by NF-κB inactivation but enhanced by NF-κB constitutive activation.

How NF-κB signaling mediates the opposite effects on intraneuronal tau inclusions triggered by exogenous tau seeds vs. those induced by transgene alone is not known. These two models develop tau inclusions with vastly different time line-1 month post-inoculation vs. 8-9 months, and likely involve distinctive proteostasis mechanisms. Beyond modulating tau phosphorylation^16^, microglial NF-κB may also modulate the sorting, trafficking and exocytosis of tau in neurons. Further investigation is needed to determine the extracellular soluble and seeding-competent tau released from microglia, and to identify the cellular machinery involved in sorting, transportation, degradation, and release of tau in both neurons and glia.

Microglia activation in neurodegenerative diseases can have both beneficial and detrimental effects. Similarly, in response to tauopathy, some of the microglial responses are adaptive, serving protective functions, while others could be maladaptive and promote toxicity. Inhibiting microglial NF-κB activity is sufficient to rescue the learning and memory deficits in PS19 mice, suggesting that NF-κB hyperactivation drives maladaptive toxic responses. In transcriptomic analyses of tau-stimulated microglia in culture and in vivo, we identified that elevation of cytokines such as IL-1, IL-6, TNFα and interferons is among the major consequences of activation of microglial NF-κB. Chronic elevation of these cytokines can cause neurotoxicity. In other models of neurodegeneration, inhibition of microglial NF-κB reduced inflammatory markers, rescued motor neuron death and extended survival of ALS mice^55^. In a kainic acid-induced neurodegenerative mouse model, deletion of IKKβ in microglia reduced expression of IL-1β and TNFα, which resulted in 30% reduction of hippocampal neuronal cell death^56^. Another potential mechanism mediating the toxic maladaptive responses could be the elevated expression of complement system and related genes (e.g. *C1qa, C1qc),* which are also positively regulated by microglial NF-κB ^57, 58^. In AD mouse models, activated microglia were found to phagocytose synapses, a process relying on the activation of complement factors, including C3, C1q and CR3^59, 60^. In tauopathy, inactivation of C3-C3aR signaling reverses the deregulation of immune network and rescues behavior deficits in PS19 mice ^61^. The exact protective mechanism underlying inactivating microglial NF-κB remains to be determined.

Our surprising finding that inactivating microglial NF-κB protected against tau-mediated cognitive deficits despite elevated tau inclusions provides compelling evidence that tau aggregates per se might not be neurotoxic. Indeed, previous studies have linked neurotoxicity with soluble tau, rather than insoluble tau aggregates ^62^. More recent studies showed that ApoE4 exacerbates tau-mediated neurodegeneration and proinflammatory responses, without increasing tau inclusions in PS19 mice^63^. We found that the protective effects of microglial NF-κB inhibition is associated with normalization of >90% of the transcriptome, and prevents the adoption of tau-mediated microglial states, in the presence of much elevated tau inclusions. Taken together, our work shows that microglial NF-κB acts downstream of tau pathology, and directly mediates toxic effects on cognition, highlighting the potential of blocking maladaptive microglial responses instead of removing tau aggregates as a therapeutic strategy to treat tauopathy.

## Methods

### Mice

P301S transgenic mice (JAX:008169) or *Cx3cr1^CreERT2/CreERT2^* mice (JAX: 021160) were crossed with *Ikbkb^F/F^* mice (MGI:2445462) or R26-Stop^FL^ikk2ca *(ikbkbCA^F/F^)* mice (JAX:008242) to obtain *P301S+;Ikbkb^F/F^, P301S+;IkbkbCA^F/F^, Cx3cr1^CreERT2/+^;Ikbkb^F/F^,* and *Cx3cr1^CreERT2/+^;IkbkbCA^F/F^* mice. *P301S+;Ikbkb^F/F^* mice were then crossed with *Cx3cr1^CreERT2/+^;Ikbkb^F/F^* mice to obtain *P301S+;Cx3cr1^CreERT2/+^; Ikbkb^F/F^* mice and littermates controls including *Ikbkb^F/F^, Cx3cr1^CreERT2/+^; Ikbkb^F/F^*, and *P301S+; Ikbkb^F/F^* mice. Similarly, *P301S+;IkbkbCA^F/F^* mice were then crossed with *Cx3cr1^CreERT2/+^; IkbkbCA^F/F^* mice to obtain *P301S+;Cx3cr1^CreERT2/+^; IkbkbCA^F/F^* mice and littermates controls including *IkbkbCA^F/F^, Cx3cr1^CreERT2/+^; IkbkbCA^F/F^*, and *P301S+; IkbkbCA^F/F^* mice. Mice were given ad libitum access to food and water and were housed in a pathogen-free barrier facility with 12-hour light on/off cycle. Mice of both sexes were used for all experiment except for RNA-seq, which used only male mice. All animal work was performed in accordance with NIH guidelines and protocols approved by University of California, San Francisco, Institutional Animal Care and Use Committee.

### Tamoxifen Administration

To induce efficient Cre expression and recombination *in vivo,* tamoxifen (T5648, Sigma-Aldrich) was dissolved in corn oil to prepare 20mg/ml stock. Mice were given tamoxifen via intraperitoneal (IP) injection at 2 mg per day for 10 days between 2.5 months to 3 months of age. To induce Cre expression and recombination in primary microglia, 4-hydroxytamoxifen (SML1666, Sigma-Aldrich) were incubated with glia mixed culture for 3 days at 5ug/ml during day 10-12 of culture. Microglia were centrifuged to completely remove 4-hydroxytamoxifen before use.

### Primary microglia culture

Primary microglia were prepared as described previously^64^. Briefly, mouse hippocampi and cortices from 2-3 day old newborn pups were isolated in DPBS. After removing the meninges, brain tissues were cut to small pieces and digested with 0.1% trypsin at 37°C for 20 min before neutralizing the trypsin with 30% FBS/DMEM media. Digested tissues were triturated to cell suspension and centrifuged for 15 min at 200g. After resuspension in 10%FBS/DMEM, cells were plated onto poly-D-lysine (PDL)-coated T-75 flasks to generate mixed glial cultures. When confluent on day 12, microglia were separated from the glia layer by shaking the flask at 200rpm for 3 hours. Floating microglia were collected for RNA analysis or seeded at 75,000 cells/cm^2^ in PDL-coated plates for tau fibril stimulation and processing assay.

### Adult microglia isolation

Adult microglia were isolated using magnetic-activated cell sorting (MACS) as described before^65^. Briefly, anesthetized mice were thoroughly trancardially perfused with cold PBS to remove circulating blood cells in the CNS. Dissected brains were chilled on ice and minced in digestion media containing with 0.2% collagenase type 3 (LS004182, Worthington) and 3 U/mL dispase (LS02104, Worthington). After 37□°C incubation for 45□min, digestion was inactivated by 2.5 mM EDTA (15575020, Thermofisher) and 1% fetal bovine serum (10082147, Thermofisher). Digested brain tissues were triturated by serological pipette to cell suspension and passed through a 70-μm cell strainer. Myelin in the cell suspension was depleted by myelin removal beads (130-096-733, Miltenyi Biotec) and magnetic LD columns (130-042-901, Miltenyi Biotec). Adult microglia were finally enriched from the eluant by CD11b MicroBeads (130-049-601, Miltenyi Biotec) and magnetic MS column (130-042-201, Miltenyi Biotec) for RNA isolation.

### RNA isolation and Quantitative PCR

To quantify target gene expression, RNA from primary microglia, isolated microglia and cortical tissue were extracted by Direct-zol RNA micro-prep kit (R2061, Zymo Research) following the manufacturer’s instructions. Genomic DNA was digested by DNase I. For qPCR, RNA was reverse-transcribed to cDNA by iScript cDNA synthesis kit (1708890, Bio-Rad). The qPCR reactions were performed using SYBR Green PCR Master Mix (4309155, Applied Biosystems) on ABI 7900HT real-time system (Applied Biosystems). GAPDH was used as a reference gene for normalization and the relative expression differences were calculated based on the 2^ΛΛCt^ method. The following primers were used for RT-qPCR: *Gapdh* (forward) 5’-GGGAAGCCCATCACCATCTT-3’, (reverse) 5’-GCCTTCTCCATGGTGGTGAA-3p; *Ikbkb*: (forward) 5’-AAGAACAGAGACCGCTGGTG-3’ and (reverse) 5’-CAGGTTCTGCATCCCCTCTG-3’

### Brain tissue collection

Mice were anesthetized with tribromoethanol and trancardially perfused with PBS. Hippocampus and cortex were dissected from one hemibrain and flash frozen at −80°C for biochemical analyses and snRNAseq, whereas the other half hemibrain (or whole brain from tau spreading experiment) was fixed in 4% paraformaldehyde for 48 h, followed by 30% sucrose infiltration for 48 h at 4°C. 30μm-thick coronal brain sections were prepared by freezing microtome (Leica) and stored at −20□ °C in cryoprotectant before staining.

### Immunohistochemistry and image analysis

DPBS was used for immunohistochemistry. 6-8 pieces brain sections per mouse that contain a series of anterior to posterior hippocampus were washed to remove cryoprotectant and then permeabilized by 0.5% Triton X-100. After blocking in 5% normal goat serum (NGS) for 1 hour, brain sections were incubated with primary antibodies in the same blocking buffer overnight at 4°C. Sections were then washed by DPBS containing 0.1% Tween-20 and incubated with Alexa-conjugated secondary antibodies for 1 hour in blocking buffer. After washing, sections were mounted on glass slides with ProLong Gold Antifade Mountant. The primary antibodies used for immunohistochemistry were as follows: anti-Iba1 (1:500, 019-19741, Fujifilm Wako), anti-MC1 (1:500, a kind gift from P. Davies). Images for MC1 and Iba1 quantification were acquired on Keyence BZ-X700 microscope using 10x objective and analyzed with ImageJ (NIH)^66^. All images were processed with auto local threshold Phansalkar plugin. Regions of interest including hippocampus and cortex were hand-traced. MC1+ areas were measured by ImageJ, whereas microglia numbers were counted with the Analyze Particles function. Images for 3D microglia reconstruction were acquired using confocal microscope LSM880 (ZEISS) at 40x magnification. Three fields per mouse of CA3 hippocampal region were taken. 3D structure of microglia was reconstructed using the Imaris software as described before^44^. Experimenters performing imaging and quantification were blinded.

### Tau fibril stimulation, uptake and clearance assay

K18/PL and full-length tau fibrils were synthesized and labeled with Alexa Fluor 647 (K18/PL) as described before^46^. All *in vitro* high content assays were performed in 96-well PDL plates (655946, Greiner) with primary microglia at a density of 25,000 cells/well. For NF-κB reporter stimulating assay, wildtype microglia were infected with Lenti-κB-dEGFP virus for 24 hours and then incubated with tau fibrils, monomers and LPS at desired concentrations for additional 24 hours. Stimulated microglia were then fixed for high content analysis. For RNA-seq analysis, wildtype microglia were plated in PDL plates for 24 hours before being stimulated with 2ug/ml full length tau fibrils for another 24 hours. RNA of microglia was then isolated for RNA-seq analysis. For tau fibril uptake assay, microglia were plated for 24 hours and then incubated with Alexa-647 labeled K18/PL fibrils (2.5ug/ml) for 6 or 24 hours. Media were changed to DMEM containing 0.01% trypsin and incubated for 5 minutes to remove tau fibrils stuck on the surface of microglia^67^. Accumulated fluorescent tau fibrils were quantified by high content assay. Lysosomes were labeled with Dextran-FITC (10,000 MW, D1820, ThermoFisher) as described before^40^. Briefly, microglia were incubated with 3mg/ml Dextran-FITC for 16 hours and then chased for at least 3 hours with fresh media before being fed with fluorescence tau fibrils. After immunostaining with Iba1, representative images were taken by confocal microscope LSM880 (ZEISS) at 60x magnification. For the clearance assay, microglia that took up tau fibrils for 24 hours were cleaned by DMEM/0.01% trypsin, followed by 10%FBS/DMEM incubation for additional 6 hours and 24 hours before harvesting for high content assay to quantify remaining fluorescent tau fibrils.

### Endotoxin Detection

Endotoxin levels in tau fibrils, monomers and LPS were detected using endotoxin detection kit following the manufacturer’s protocol (GenScript ToxinSensor™ Chromogenic LAL Endotoxin Assay Kit). All tau samples had endotoxin levels of <1.0 EU/mL at working concentration.

### High content analysis assay

High content analysis assay, including immunostaining, images acquiring and automated analysis, was used to unbiasedly detect and quantify immunofluorescence signals in cultured cells^68^. After treatment and trypsin cleaning, microglia were fixed with conditioned medium containing 4% paraformaldehyde, permeabilized with 0.1%Triton X-100, and incubated for 1 hr in blocking solution containing DPBS, 0.01% Triton X-100, and 5% NGS. The cells were then incubated in blocking solution containing anti-Iba1 antibody overnight at 4°C, followed by incubation with secondary antibody for 1 hr. Nuclei were labeled with Hoechst (H3570, Invitrogen). Images from individual wells were acquired with a fully automated ArraySan XTi high-content system (Thermo Fisher) and a 20x objective. Twenty-five fields containing 3,000-5,000 microglia and encompassing the total well area were captured. Images were auto-quantified with CellHealthProfile module of Cellomics software (Thermo Fisher). Iba1 signal was used to define the quantification area and count microglia number. Total intensity of dEGFP or fluorescence tau fibrils within the Iba1 area were quantified and normalized to microglia number in each well.

### High-throughput bulk RNA sequencing

The following sample sets were prepared for bulk RNA-seq analysis. Set 1 (related to Fig.1a,b,g,i, supplementary Fig.1c): N=4 replicates of wildtype mouse primary microglia treated with vehicle or 0.5 μg/ml full length 0N4R tau fibrils for 24 hours; Set 2 (related to Fig.1e,f,h,j, supplementary Fig. 1d): N=4 replicates of adult microglia isolated from 11-month-old male non-transgenic control mice and PS19 mice; Set 3 (related to Fig. 2c-e,h-j, supplementary Fig. 2): N=5 replicates of primary microglia with endogenous IKKβ expression or IKKβCA overexpression and N=4 replicates of primary microglia with wildtype IKKβ expression or deletion of IKKβ; Set 4 (related to Fig. 6): N=4 cortical tissues from *Ikbkb^+/+^, Ikbkb^-/-^, Ikbkb^+/+^;P301S+ and Ikbkb^-/-^;P301S+* male mice. For all sample sets, total RNA was extracted using Quick-RNA miniprep (R1055, Zymo Research). RNA quality was examined by a 2100 Bioanalyzer Instrument (Agilent Genomics). RNA samples with RNA integrity numbers greater than 8 were used for complementary DNA library construction. For sample set 1-3, cDNA library generation was done using the QuantSeq 3 mRNA-Seq Library Prep Kit (FWD) for Illumina (K01596, Lexogen) following manufacturer’s instructions. The qualities of the cDNA library were assessed using Qubit 2.0 fluorometer to determine the concentrations and 2100 Bioanalyzer Instrument to determine insert size. cDNA library samples were then sequenced with the HiSeq 4000 System (Illumina). For sample set 4, cDNA library generation and RNA-seq service was performed by Novogene (Novogene Co., Ltd, Sacramento, California). Briefly, oligo(dT) beads were used to enrich for mRNA. After chemical fragmentation, the cDNA libraries were generated using NEBNext Ultra RNA Library Prep Kit for Illumina (E7530S, New England Biolabs). Quality of the cDNA library was also assessed by Qubit assay for concentration, Bioanalyzer 2100 for insert size, and qPCR for effective library size. cDNA library samples were then sequenced with the HiSeq 4000 at PE150. On average, 0.02-0.03% error rate was detected. Raw reads were filtered by (1) remove reads containing adaptors; (2) remove reads containing N > 10% (N represents base that could not be determined); (3) The Qscore (Quality value) of over 50% bases of the read is <= 5.

### Data analyses of bulk RNA-seq

For sample set 1-3, RNA-seq reads were mapped using the BlueBee genomics platform and the GENCODE GRCm38 mouse genome (Lexogen QuantSeq 2.2.3) as reference genome. For sample set 4 and 5, RNA-seq read mapping was performed using the STAR program^69^ and same GENCODE GRCm38 mouse genome was used as reference. Total mapping rate is more than 95% and unique mapping rate is >90%. The read count table was generated with the RSEM program^70^. Differential gene expression was calculated with R package edgeR^71^ and limma^72^. Counts were normalized with the trimmed mean normalization method^73^. FDR was calculated using the Benjamini-Hochberg method. DE genes were defined with FDR less than 0.05 in comparison to control group. Gene network analyses of RNA-seq data were performed using Ingenuity Pathway Analysis (IPA, Qiagen).

### Isolation of nuclei from frozen mouse brain tissue

The protocol for isolating nuclei from frozen postmortem brain tissue was adapted from a previous study with modifications^48, 74^. All procedures were done on ice or at 4°C. In brief, mouse brain tissue was placed in 1500 μl of Sigma nuclei PURE lysis buffer (Sigma, NUC201-1KT) and homogenized with a Dounce tissue grinder (Sigma, D8938-1SET) with 15 strokes with pestle A and 15 strokes with pestle B. The homogenized tissue was filtered through a 35-μm cell strainer and were centrifuged at 600 g for 5 min at 4°C and washed three times with 1 ml of PBS containing 1% BSA, 20 mM DTT and 0.2 U μl^-1^ recombinant RNase inhibitor. Then the nuclei were centrifuged at 600 g for 5 min at 4°C and re-suspended in 500 μl of PBS containing 0.04% BSA and 1x DAPI, followed by FACS sorting to remove cell debris. The FACS-sorted suspension of DAPI-stained nuclei were counted and diluted to a concentration of 1000 nuclei per microliter in PBS containing 0.04% BSA.

### Droplet-based single-nucleus RNA sequencing

For droplet-based snRNA-seq, libraries were prepared with Chromium Single Cell 3’ Reagent Kits v3 (10x Genomics, PN-1000075) according to the manufacturer’s protocol. cDNA and library fragment analysis was performed using the Agilent Fragment Analyzer systems. The snRNA-seq libraries were sequenced on the NovaSeq 6000 sequencer (Illumina) with 100 cycles.

### Analysis of Droplet-Based Single-nuclei RNA-seq

Gene counts were obtained by aligning reads to the mouse genome (mm 10) with Cell Ranger software (v.3.1.0) (10x Genomics). To account for unspliced nuclear transcripts, reads mapping to pre-mRNA were counted. Cell Ranger 3.1.0 default parameters were used to call cell barcodes. We further removed genes expressed in no more than 2 cells, cells with unique gene counts over 8,000 or less than 300, and cells with high fraction of mitochondrial reads (> 5%). Potential doublet cells were predicted and removed using DoubletFinder ^49^ for each sample. Normalization and clustering were done with the Seurat package v3.0.1 ^75^. In brief, counts for all nuclei were scaled by the total library size multiplied by a scale factor (10,000), and transformed to log space. A set of 2000 highly variable genes were identified with FindVariableFeatures function based on a variance stabilizing transformation (vst). Principal component analysis (PCA) was done on all genes, and t-SNE was run on the top 15 PCs. Cell clusters were identified with the Seurat functions FindNeighbors (using the top 15 PCs) and FindClusters (resolution = 0.1). For each cluster, we assigned a cell-type label using statistical enrichment for sets of marker genes and manual evaluation of gene expression for small sets of known marker genes. The subset() function from Seurat was used to subset microglia-only cells. Differential gene expression analysis was done using the FindMarkers function and MAST ^76^.

### Cell trajectory using Monocle3

For microglial trajectory analysis, the microglial population was first isolated from the other cell types using the previously identified cell types. A separate Seurat object was created for microglia, followed by normalization with a scale factor of 10,000. FindVariableFeatures function was run again to identify the most variable gene specific for microglia. The microglial Seurat object was then converted into a Monocle3 object with as.cell_data_set function. Size factor estimation of the new CDS (cell data set) was performed using estimate_size_factors function with default parameters. Further processing of the CDS was carried out using preprocess_cds function with the num_dim parameter set to 9. UMAP was then performed to reduce the dimensionality of the data. Cell clusters were then visualized with cluster_cells function with the parameter K equals to 9. learn_graph function was used to determine the trajectory and the *Ikbkb^-/-^;P301S+* cluster was selected as the origin of the trajectory.

### Gene network and functional enrichment analysis

Gene network and functional enrichment analysis were performed by Ingenuity Pathway Analysis (IPA, Qiagen) or by GSEA with molecular signatures database (MSigDB)^77, 78^. Significant DEGs and their log2 fold change expression values and FDR were inputted into IPA for identifying canonical pathways, biological functions and upstream regulators. Upregulated or downregulated significant DEGs were inputted into GSEA to identify hallmark and gene ontology pathways. The p-value, calculated with the Fischer’s exact test with a statistical threshold of 0.05, reflects the likelihood that the association between a set of genes in the dataset and a related biological function is significant. A positive or negative regulation z-score value indicates that a function is predicted to be activated or inhibited.

### Plexxikon drug administration

Plexxikon diet containing 1200 mg/kg PLX5622 compound (Plexxikon Inc., Berkeley) was used as the sole food source for 2 weeks to deplete microglia before inoculation of tau seeds. Control diet with the same base formula but no compound was used for control group. Mice were fed with plexxikon diet throughout the whole experiment to suppress microglial repopulation^44^.

### Preparation of PSP brain extract

Human brain tissues from PSP cases, which were diagnosed based on accepted neuropathology criteria, were obtained from Dr. Lea Grinberg at UCSF Memory and Aging Center. The use of postmortem brain tissues for research was approved by the University of California, San Francisco’s Institutional Review Board with informed consent from patients or their families. Brain extracts were prepared as previously described^79^. Briefly, PSP brain tissue was homogenized at 10% (w/v) in sterile phosphate-buffered saline (PBS) with protease, phosphotase and HDAC inhibitors. After brief sonication (power 40, 5min, 10 sec cycles), the homogenates were centrifuged at 3000 g for 5 min at 4 °C. The protein concentration of the supernatant was measured by BCA assay and then aliquoted and frozen at −80°C until use.

### Stereotaxic injection

The stereotaxic surgery was performed as previously described^80^. Briefly, 3 month-old mice that had been fed with PLX or control diet or injected with tamoxifen were anesthetized with 2% isoflurane by inhalation during the surgery. When deeply anaesthetized, mice were secured on a stereotaxic frame (Kopf Instruments) and were unilaterally injected to the right hippocampus using CA1 coordinates (bregma, −2.5 mm; lateral, +2 mm; and depth, −1.8 mm) with a Hamilton syringe under aseptic conditions. The following tau seeds were inoculated: (1) 3 μL of 4.3 μg/ul PSP brain extracts for microglia-depleted PS19 mice (Supplementary Fig. 4); (2) 2 μl of 2.5 μg/ul K18/PL tau fibrils for microglia-depleted PS19 mice (Fig. 4a-c) and PS19 mice with microglial NF-κB inactivation (Fig.4 f-h); (3) 2 μL of 0.2 μg/ul K18/PL tau fibrils for PS19 mice with microglial NF-κB activation (Fig.4 k-m). Same volumes of PBS were injected to PS19 mice or same amount of tau seeds were injected to non-transgenic mice as control. Mice were monitored during the anesthesia until recovery. After 3 months (for PSP brain extracts) or 1 month (for K18/PL tau fibrils) of spreading, mice were perfused for whole brain immunohistochemistry of AT8 or MC1 tau as described.

### Morris water maze

Morris water maze studies were conducted during daylight hours. The water maze consisted of a pool (122 cm in diameter) containing opaque water (20 ± 1°C) and a platform (14 cm in diameter) submerged 1.5 cm below the surface. Hidden platform training (days 1-6) consisted of 12 sessions (two per day, 2 h apart), each with two trials. The mouse was placed into the pool at alternating drop locations for each trial. A trial ended when the mouse located the platform and remained motionless on the platform for 5 s, for a maximum of 60 s per trial. Mice that failed to find the platform within the 60 s trial were led to it and placed on it for 15 s. Probe trails were conducted 72 hours after the final hidden training. Mice were returned to the pool with a drop location that was 180° opposite of original target platform location in the absence of the hidden platform. Performance was measured with EthoVision video-tracking (Noldus Information Technology). Visible platform training, where the platform was cued with a mounted black-and-white striped mast, was conducted for three sessions after the conclusion of probe trials. Pre-set criteria for exclusion from analysis included floating and thigmotaxic behaviors, neither of which was observed in current studies.

### Statistics

The sample size for each experiment was determined on the basis of previous experience. All *in vitro* experiments were performed with a minimum of 3 biological replicates. Mean values from at least three independent experiments were used for computing statistical differences. All *in vivo* experiments were performed with a minimum of 4 mice per genotype. All *in vivo* data were averaged to either individual mouse (microglia number counts), individual section (MC1, AT8 tau) or individual microglia (Imaris morphology analysis) and mean values were used for computing statistical differences. Statistical analyses were performed with Graphpad prism 8.0 (Graphpad, San Diego, California), STATA 12 (StataCorp) and R (R Foundation for Statistical Computing, Vienna, Austria). Data visualization were done with Graphpad and R package ggplot2^81^. Values are reported as mean ± standard error of the mean (SEM). The Shapiro-Wilk test of normality and F test to compare variances were applied to data sets when applicable. Unpaired *t* test was used to compare 2 groups. One-way ANOVA was used to compare data with more than 2 groups. Two-way ANOVA was used for groups with different genotypes and/or time as factors. Tukey’s and Sidak’s post-test multiple comparisons was used to compare statistical difference between designated groups. Multilevel mixed-effects linear regression model was used when parameters vary at more than one level. *P*□<□0.05 was considered statistically significant

## Acknowledgements

The authors thank Dr. Peter Davies from Albert Einstein College of Medicine for MC1 antibody; Dr. Michael Karin at University of California-San Diego and Dr. Katerina Akassoglou at Gladstone Institutes for the *Ikbkb^F/F^* mice; Dr. Michael Gill at the Gladstone Behavior Core for the MWM test assistance; Dr. Meredith Calvert at the Gladstone Histology and Light Microscopy Core for imaging assistance; Dr. Anke Meyer-Franke at the Gladstone Assay Development and Drug Discovery Core for the high content assay assistance; Dr. Santiago Sole Domenech and Dr. Fred Maxfield at Weill Cornell Medicine for lysosome tracking; the University of California, San Francisco’s Center for Advanced Technology for conducting bulk RNA sequencing; Jason McCormick and Tomas Baumgartner from Weill Cornell Medicine Flow Cytometry Core Facility for FACS assistance; Dong Xu, Xing Wang and Adrian Tan from Genomics Resources Core Facility for performing single nuclei RNA sequencing. **Funding:** This work was supported in part by the National Institute of Health Grants R01AG051390, U54NS100717, R01AG054214, and Rainwater Foundation (to L.G), the National Institute of Aging Grant F30 AG062043-02 and National Institute of Health Grant T32GM007618 (to L.K.).

## Author contributions

C.W. and L.G. designed research; C.W., L.F., L.Z., L.K., M.C.,Y.L., D.L., Y.Z., C.C. performed experiments; S.M., J.G. contributed reagents preparation; C.W., L.F., L.Z., L.K., M.C., B.L., Y.L., D.L., C.C., L.G. analyzed data; C.W., L.F., B.L. and L.G. wrote the manuscript.

## Supplementary Information

**Supplementary Fig. 1.**
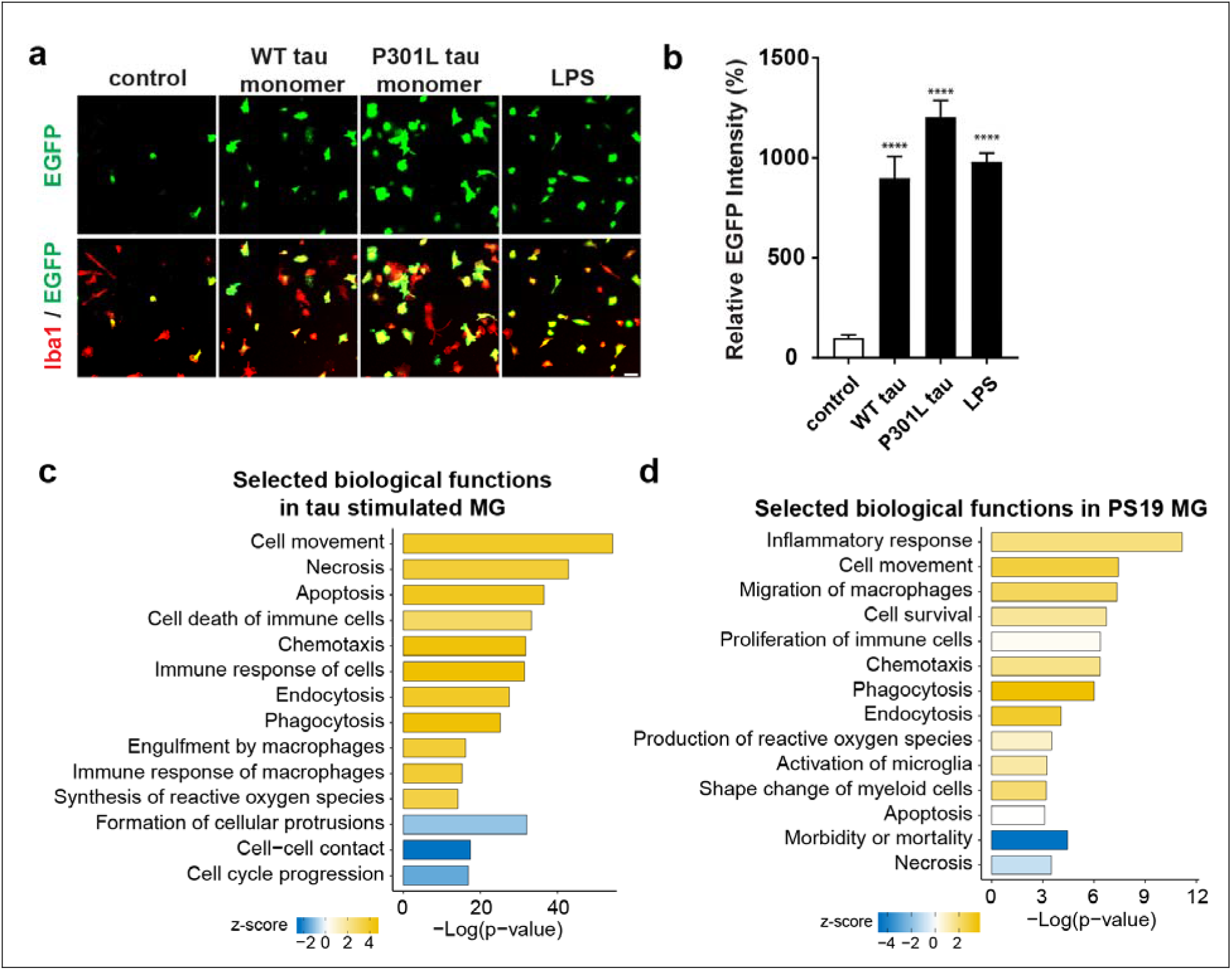
(related to Fig. 1) Tau activates NF-κB pathway and modulates biological functions in microglia. **(a,b)** Primary microglia infected with NF-κB reporter (Lenti-κB-dEGFP) virus were incubated with wildtype (100nM), P301L (100nM) tau monomers and LPS (50ng/ml) for 24h. (**a**) Representative fluorescence high content images of EGFP (green) and Iba1 (red); (**b**) Quantification of EGFP intensity. Scale bar, 50μm. Data are from two independent experiments, total N=8 wells. Values are mean ± SEM, relative to vehicle control, one-way ANOVA with Tukey’s multiple comparison posttest, ****p<0.0001 **(c,d)** Selected IPA biological functions identified for DEGs in tau stimulated (**c**) or PS19 (**d**) microglia.

**Supplementary Fig. 2.**
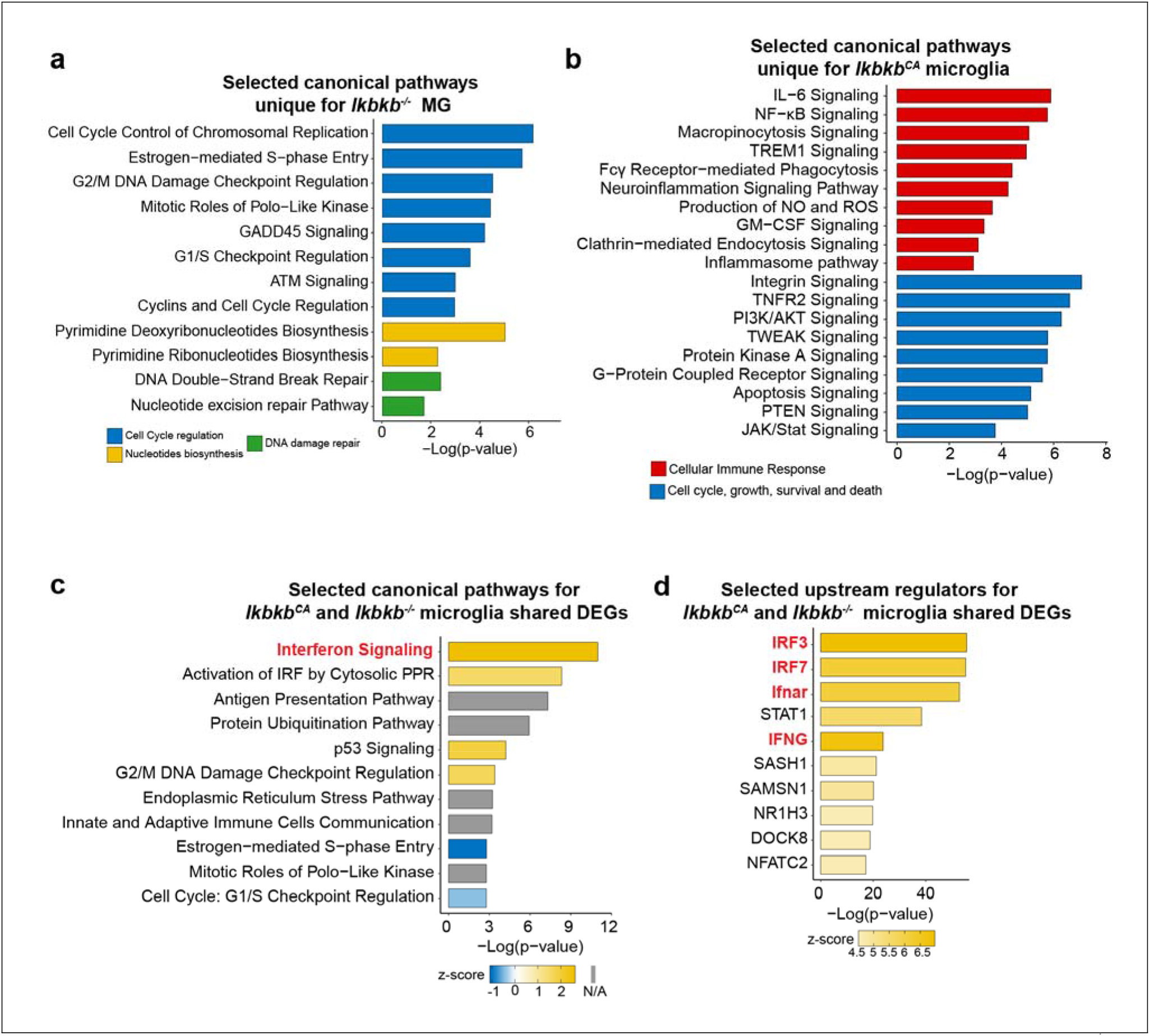
(related to Fig. 2) IPA analysis of unique and shared DEGs of *Ikbkb^-/-^* and *Ikbkb^CA^* microglia. **(a,b)** Selected IPA canonical pathways identified for unique DEGs of *Ikbkb^-/-^* microglia (**a**) and *Ikbkb^CA^* microglia (**b**). Canonical pathways are grouped by indicated categories. **(c,d)** Selected IPA canonical pathways (**c**) and upstream regulators (**d**) identified for shared DEGs of *Ikbkb^-/-^* and *Ikbkb^CA^* microglia.

**Supplementary Fig. 3.**
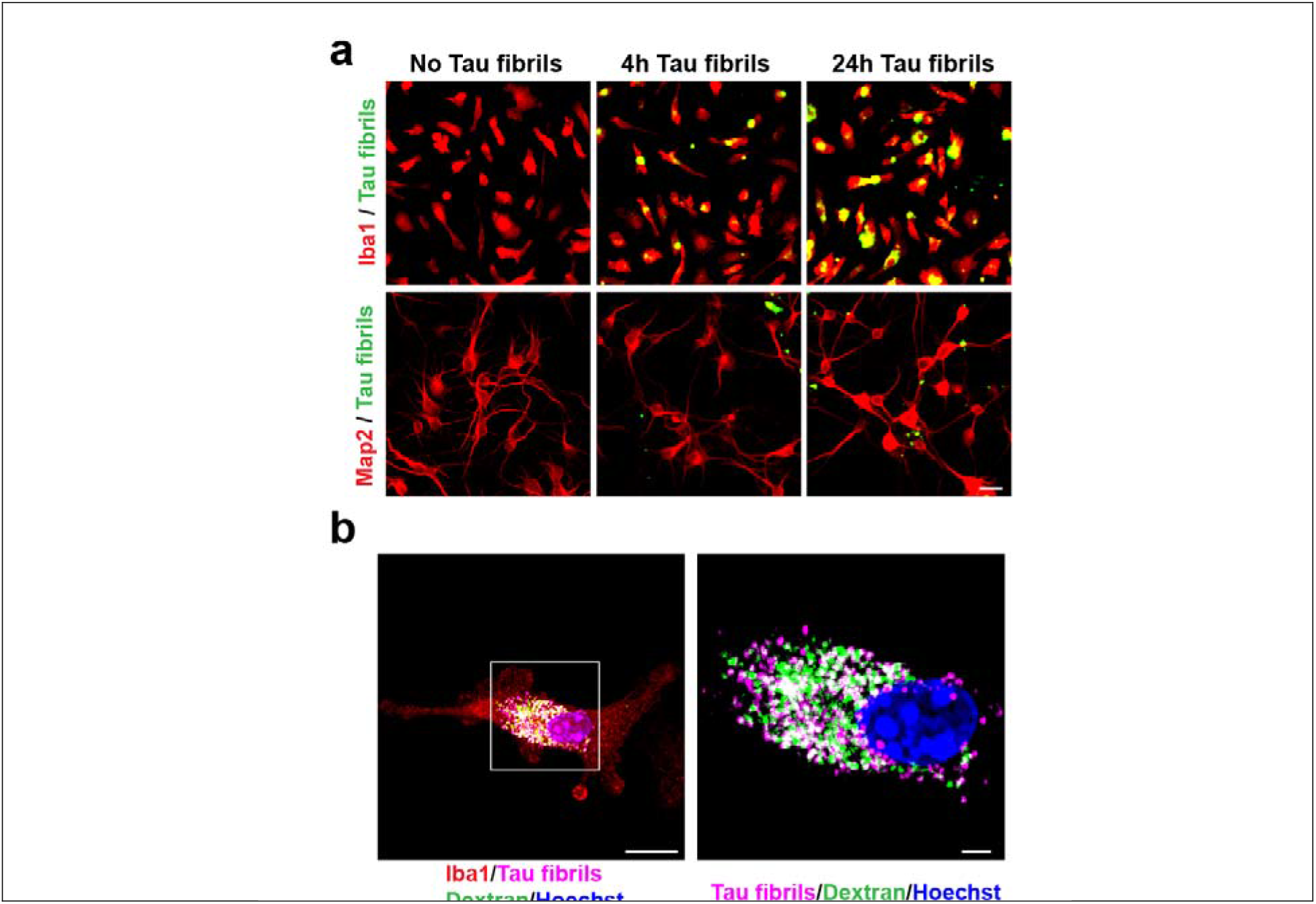
(related to Fig. 3) Microglia take up tau fibrils. **(a)** Primary microglia (Iba1) and neurons (Map2) were incubated with fluorescent tau fibrils for 4 and 24 hours. Representative images show that microglia, but not neurons, take up tau fibrils in a time-dependent manner. Scale bar, 25 μm. **(b)** Representative images of immunocytochemical staining show that tau fibrils were colocalized with lysosomes in microglia. Lysosomes were traced by Dextran-FITC, followed by incubation of fluorescent tau fibrils for 3 hours. Nuclei were labeled by Hoechst. Scale bar, left 10 μm, right 2 μm.

**Supplementary Fig. 4.**
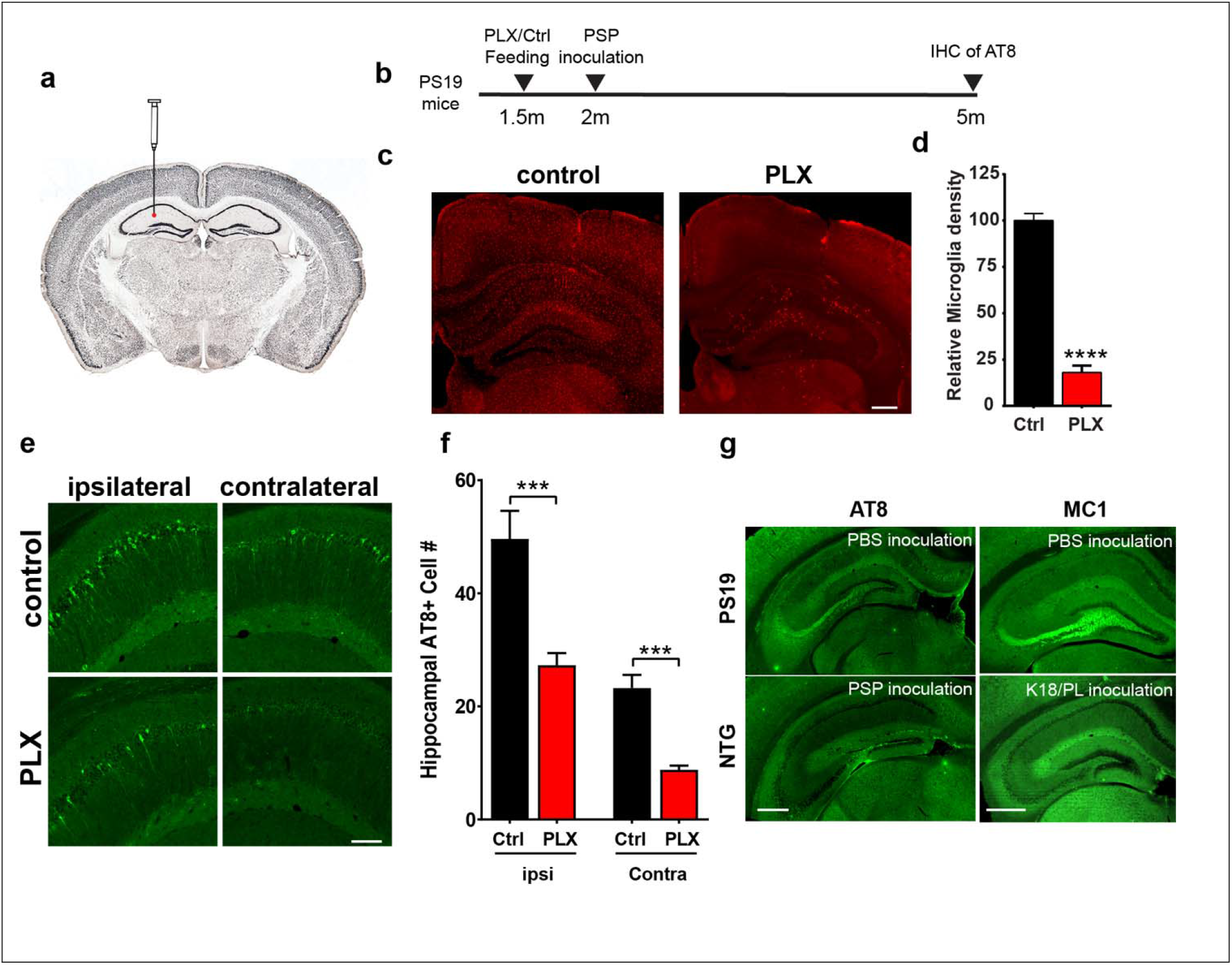
(related to Fig. 4) Depletion of microglia in PS19 mice halts tau seeding and spreading. **(a)** Schematic diagram illustrating tau seeds inoculation path. The red dot indicates injection site in hippocampus. **(b)** Schematic diagram illustrating the experimental design and timeline of microglia depletion and PSP brain extract inoculation in PS19 mice. Pathological tau seeding and spreading was determined by immunohistochemical staining of AT8. **(c,d)** Representative images (**c**) and quantification (**d**) of Iba1 positive microglia in control diet fed mice (n=6) and PLX diet fed mice (n=8). ****p<0.0001, student unpaired t-test. Scale bar, 500 μm **(e,f)** Representative immunohistochemical staining of AT8 tau in ipsilateral and contralateral hippocampus CA1 region (**e**) and quantification of AT8+ neurons number (**f**) in control diet fed mice (n=6) and PLX diet fed mice (n=9), 3 sections per mice. *** p<0.001, STATA mixed model. Scale bar, 200μm **(g)** Representative immunohistochemical staining of AT8 in PS19 mice inoculated with PBS and in non-transgenic mice inoculated with PSP brain extract (left) and MC1 in PS19 mice inoculated with PBS and in non-transgenic mice inoculated with K18/PL tau fibrils (right). Scale bar, 500μm

**Supplementary Fig. 5.**
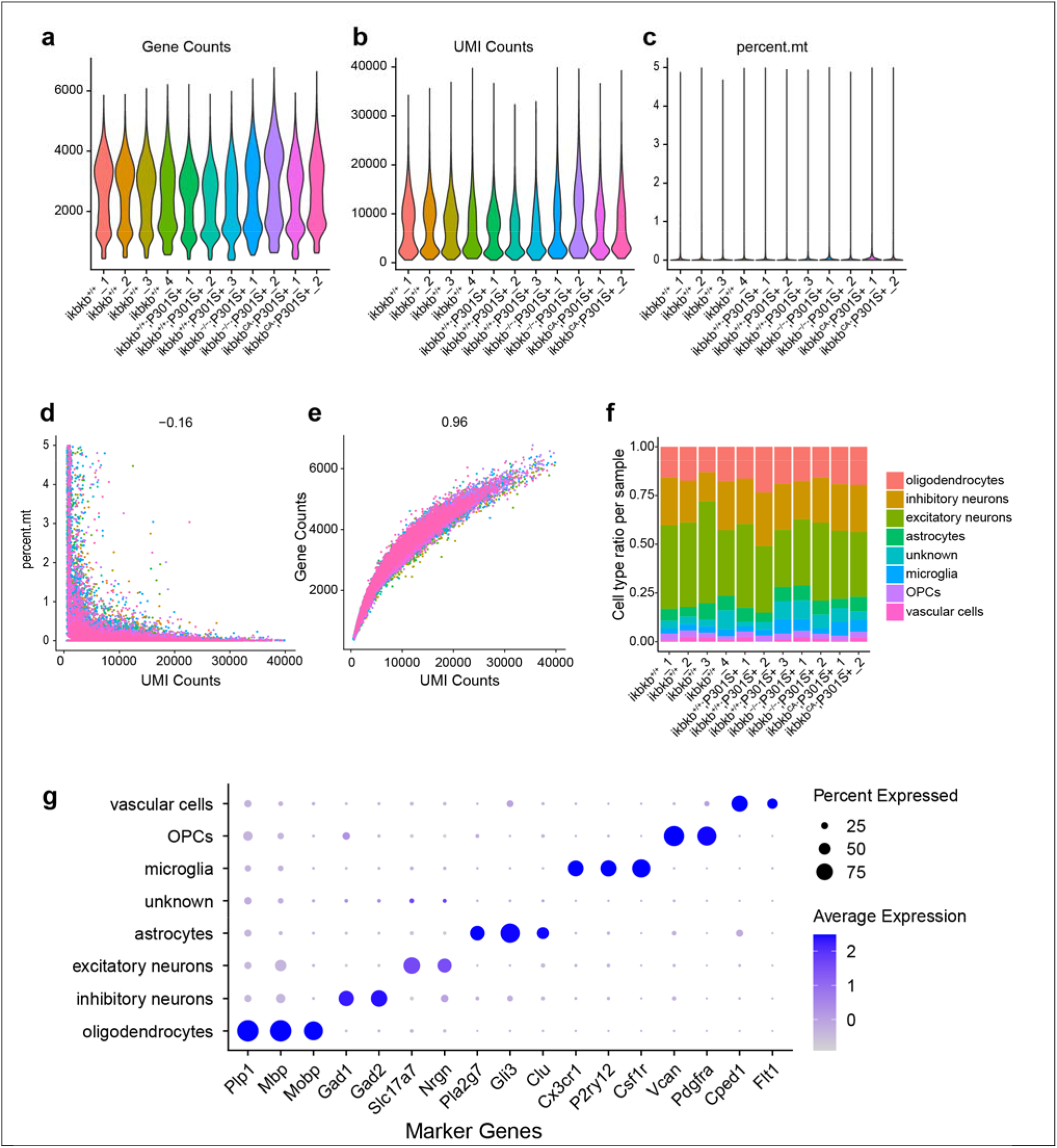
(related to Fig. 7) Quality control assessment of single nuclei RNA-Seq of cortical tissues. **(a-c)** Violin plots showing spread of total genes (**a**), total UMIs (**b**), and percent of mitochondrial genes (**c**) detected per nuclei for each individual sample. (**d,e**) Correlation between UMI counts and percentage of mitochondrial genes per nuclei (**d**) and total genes detected (**e**) for all samples. **(f)** Proportion of cell types for each individual sample. **(g)** Percentage and average expression levels of maker genes for indicated cell types.

**Supplementary Fig. 6.**
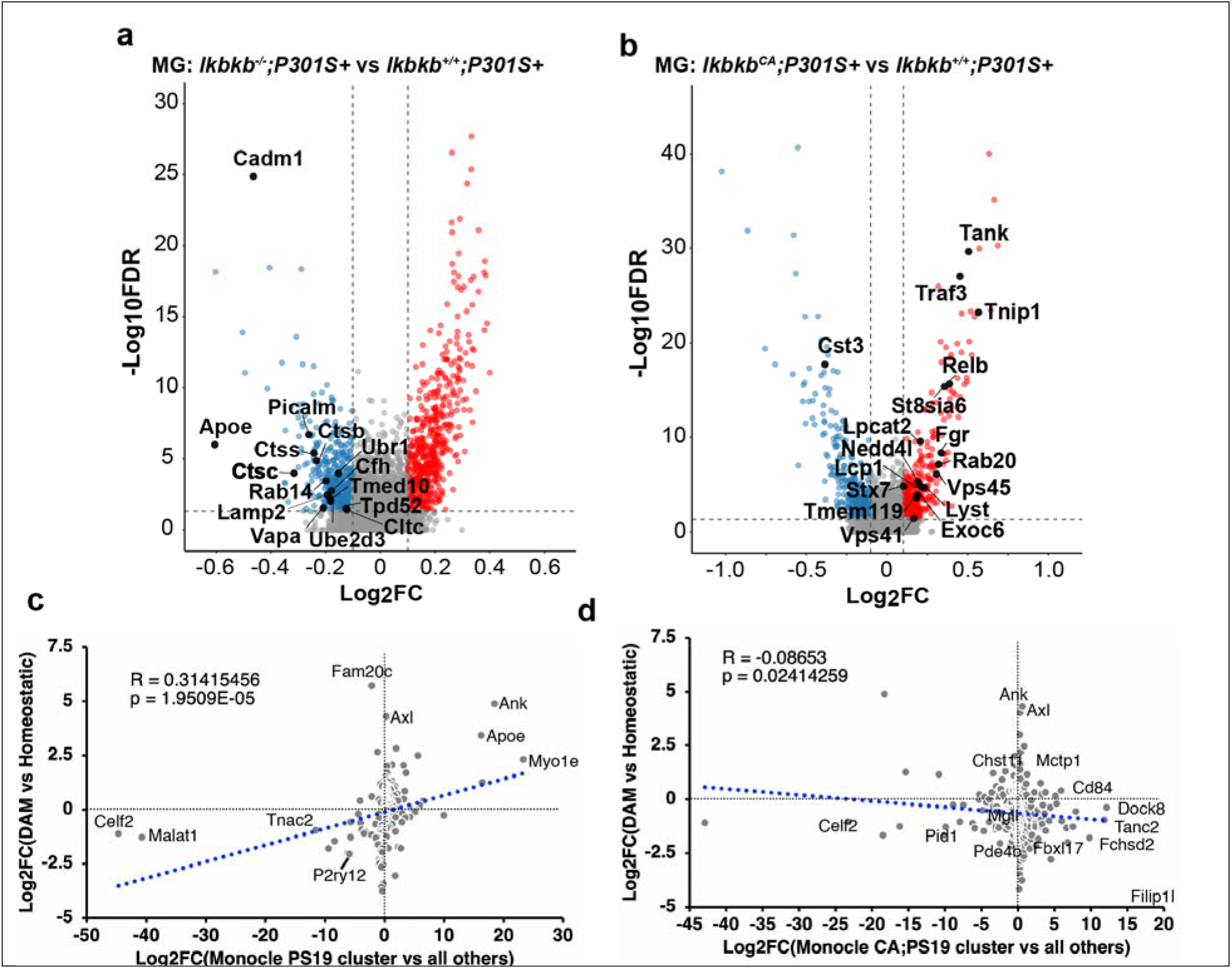
(related to Fig. 7) DEGs regulated by microglial NF-κB in PS19 mice. **(a,b)** Volcano plots of significant DEGs (FDR<0.05) from microglia in *Ikbkb^-/-^;P301S+* mice (**a**) and *Ikbkb^CA^;P301S+* mice (**b**), in comparison to microglia from *Ikbkb^+/+^;P301S+* mice. Selected DEGs related GESA pathways are labeled. (**c**) Correlation of genes markers of cluster 3 vs. all other clusters with DAM genes vs. homeostatic genes. (**d**) Correlation of gene markers of cluster 4 and 5 vs. all other clusters with DAMs genes vs. homeostatic genes.

