## Supplementary table_1 for "Microglial NF-κB drives tau spreading and toxicity in a mouse model of tauopathy"

Supplementary Table 1. Endotoxin levels of tau monomer and fibrils

| **Sample** | **Conc.** | **Endotoxin (EU/ml)** |
| --- | --- | --- |
| monomer tau WT | 100nM | 0.13 |
| monomer tau P301L | 100nM | 0.16 |
| fibrillar tau K18/PL | 2.5μg/ml | 0.1 |
| fibrillar tau 0N4R | 2μg/ml | 0.88 |
| LPS | 50ng/ml | 226.0 |
